# Mapping the Psychosis Spectrum – Imaging Neurosubtypes from Multi-Scale Functional Network Connectivity

**DOI:** 10.1101/2025.02.11.637551

**Authors:** Ram Ballem, Pablo Andrés-Camazón, Kyle M. Jensen, Prerana Bajracharya, Covadonga M. Díaz-Caneja, Juan R Bustillo, Jessica A. Turner, Zening Fu, Jiayu Chen, Vince D. Calhoun, Armin Iraji

**Author notes:** Corresponding Author: Ram Ballem, Postal Address: 55 Park Place NE 18^th^ Floor – TReNDS Center Atlanta, GA – 30303, Name: Armin Iraji, Postal Address: 55 Park Place NE 18^th^ Floor – TReNDS Center Atlanta, GA – 30303.

## Abstract

This study aims to identify Psychosis Imaging Neurosubtypes (PINs)— homogeneous subgroups of individuals with psychosis characterized by distinct neurobiology derived from imaging features. Specifically, we utilized resting-state fMRI data from 2103 B-SNIP 1&2 participants (1127 with psychosis, 350 relatives, 626 controls) to compute subject-specific multiscale functional network connectivity (msFNC). We then derived a low-dimensional neurobiological subspace, termed Latent Network Connectivity (LNC), which captured system-wide interconnected multiscale information across three components (cognitive-related, typical, psychosis-related). Projections of psychosis participants’ msFNC onto this subspace revealed three PINs through unsupervised learning, each with distinct cognitive, clinical, and connectivity profiles, spanning all DSM diagnoses (Schizophrenia, Bipolar, Schizoaffective). PIN-1, the most cognitively impaired, showed Cerebellar-Subcortical and Visual-Sensorimotor hypoconnectivity, alongside Visual-Subcortical hyperconnectivity. Most cognitively preserved PIN-2 showed Visual-Subcortical, Subcortical-Sensorimotor, and Subcortical-Higher Cognition hypoconnectivity. PIN-3 exhibited intermediate cognitive function, showing Cerebellar-Subcortical hypoconnectivity alongside Cerebellar-Sensorimotor and Subcortical-Sensorimotor hyperconnectivity. Notably, 55% of relatives aligned with the same neurosubtype as their affected family members—a significantly higher rate than random chance (p-value_Relatives-to-PIN-1_ < 0.001, p-value_Relatives-to-PIN-2_ < 0.05, p-value_Relatives-to-PIN-3_ < 0.001) compared to a non-significant 37% DSM-based classification, supporting a biological basis of these neurosubtypes. Cognitive performance reliably aligns with distinct brain connectivity patterns, which are also evident in relatives, supporting their construct validity. Our PINs differed from original B-SNIP Biotypes, which were determined from electrophysiological, cognitive, and oculomotor data. These findings underscore the limitations of DSM-based classifications in capturing the biological complexity of psychotic disorders and highlight the potential of imaging-based neurosubtypes to enhance our understanding of the psychosis spectrum.

## INTRODUCTION

### 1.1. Neurobiology of Psychosis, Diagnosis and Treatment

Psychosis is believed to be a manifestation of disordered activity within the human brain, affecting thought and behavior, and impairing social and occupational functioning (1). Disorders like Schizophrenia, Bipolar Disorder with Psychosis, and Schizoaffective Disorder are complex and biologically heterogenous, with overlapping symptoms (2,3).

The diagnosis of psychotic disorders relies on DSM-5 criteria, which categorizes individuals based on comprehensive assessment of symptom patterns, co-occurring with distress and/or functional impairment. However, it ignores underlying heterogeneous biology. Despite ongoing efforts to comprehend the etio-pathophysiology of psychosis (1,4), neurobiological complexity continues to hinder the development of effective treatments (5).

Previous studies addressing the heterogeneity of psychotic disorders have focused on identifying homogenous biotypes based on cognitive, clinical, or neurophysiological characteristics (6–8). These findings suggest that subgrouping, beyond traditional DSM boundaries, can yield biologically meaningful insights, enhancing diagnosis and inspiring frameworks (9) that guide the development of targeted treatments (10).

### 1.2. Neuroimaging Insight into Psychosis

Neuroimaging has revealed aberrant brain circuitry associated with psychotic symptoms (11–13). Resting-state fMRI (rsfMRI) is a powerful non-invasive method that relates to neural activity in the brain, measured through blood-oxygenation-level-dependent (BOLD) signals (14,15), effectively identifying abnormal brain function in various neurological and psychiatric conditions including psychosis, trauma, and Alzheimer’s disease (16–22). When combined with machine learning, fMRI holds great promise for transdiagnostic research by identifying precise biomarkers that can enhance clinical outcomes across multiple disorders (23).

Spatial independent component analysis (ICA) has identified intrinsic connectivity networks (ICNs) (24) - groups of brain regions that work together. Studies using ICA in large, case-control studies have found that individuals with psychosis often show abnormal brain activity in ICNs (25,11,26,27). These abnormalities involve aberrant connectivity in the default mode network, frontoparietal (cognitive control or central executive) (28–30), and salience network (31,32), which composes the triple network model described by (33). While rsfMRI studies highlight these differences in brain activity, there is considerable variability among individuals with psychosis (34), thus suggesting that relying solely on conventional group analysis methods is insufficient to advance clinical translation of neuroscientific findings (35).

rsfMRI has been valuable in informing psychosis subgroups with distinct brain activation and connectivity patterns linked to cognitive and clinical variables (16,36). These biotypes could be more homogeneous in therapeutic response and etiology (5), aiding in overcoming challenges in psychosis research and enhancing treatment development (8). Expanding rsfMRI studies to include comprehensive brain connectivity could enhance our understanding, complement existing research, and better define a clear homogenous neurobiological boundary to uncover core variations that remain unknown.

### 1.3. Data-Driven Harmonization Across Cohorts through NeuroMark

rsfMRI data-driven approaches, such as Multivariate Objective Optimization ICA with reference (MOO-ICAR) (37), enhance conventional approaches by incorporating spatial templates as references to guide decomposition into functional brain networks. These techniques enhance standardization, accounts for subject variability, and enable reliable cross-study comparisons, ultimately improving replicability of findings (38,39). Their effectiveness depends on a robust, generalizable reference derived from large, population-representative samples.

The Neuromark 2.2 framework (38,40) advances these efforts, providing a comprehensive set of multi-spatial-scale ICNs (msICNs) based on rsfMRI data from over 100,000 individuals. Multispatial scale (or multiscale) refers to analyzing brain networks across various levels of spatial granularity - from large-scale to granular networks, enabling detailed insights into functional brain organization (38,39), and providing a wider understanding of connectivity patterns across different spatial scales.

Standardized frameworks like Neuromark 2.2, address challenges in fMRI research, by maintaining consistent comparisons across cohorts and disorders while improving reproducibility (5,38). Multiscale functional network connectivity (msFNC) derived from activations within and between spatially distinct networks, has proven crucial for understanding neural mechanisms in psychosis, addressing brain heterogeneity, and enhancing the replicability of findings (16).

### 1.4. Latent Space of Functional Network Connectivity

The human brain operates as a complex, interconnected system, where functional elements collaborate to process and respond to stimuli (41,42). However, the mechanisms driving these interactions - interconnected system-wide activity - remain poorly understood, posing a major challenge.

A promising approach to address this challenge is leveraging dimensionality reduction techniques to exploit redundancy to obtain critical coordinated patterns within complex systems (42). For instance, a recent study on stroke patients identified brain connectivity disruptions through low dimensional embeddings, identifying accurate diagnosis severity and recovery prediction (43).

Building on this concept, our objective is to investigate psychosis heterogeneity through brain activity within a low-dimensional neurobiological subspace. This focuses on examining brain-wide connectivity and its interconnected patterns, spanning large-scale network activity and detailed granular network interactions. We hypothesize that this subspace, enriched with multiscale connectivity, can uncover latent brain interaction patterns and identify clinically meaningful homogenous subgroups within the psychosis spectrum based on neurobiological features, which we define as Psychosis Imaging Neurosubtypes (PINs) (44).

## MATERIALS AND METHODS

### 2.1. Participants

The imaging data comprises rsfMRI from 2,103 participants recruited by the Bipolar-Schizophrenia Network on Intermediate Phenotypes (B-SNIP 1 & 2) Consortium (45), collected from multiple sites at participating institutions. Participants with psychosis met the criteria for Bipolar I Disorder (BP), Schizophrenia (SZ), or Schizoaffective Disorder (SAD), using the Structured Clinical Interview (SCID) (46). First-degree relatives of psychosis participants included parents, siblings, or children. Controls did not have a history of psychosis syndromes or recurrent mood syndromes. Participants were assessed with multiple cognitive and clinical scales (details in Supplementary Section 1).

Cognitive assessments were administered through Brief Assessment of Cognition in Schizophrenia (BACS) (47,48). These involved - Verbal Memory (BACS-VM), Digit Sequencing (BACS-DS), Token Motor (BACS-TM), Verbal Fluency (BACS-VF), Symbol Coding (BACS-SC), Tower of London (BACS-ToL), with a composite score (BACS-COMP) summarizing overall performance. Additional measures included Wechsler Memory Scale (49) subtests, capturing forward (WMS-F) and backward (WMS-B) subtests. Both BACS and WMS scores are normalized and stratified across age and sex. Normative values regarding this transformation are noted here (48).

Clinical characteristics rated for both controls and patients included - total scores of Birchwood Social Functioning Scale (BSFS) (50) and Global Assessment of Functioning (GAF) (51). Psychosis was rated on total score of Montgomery-Asberg Depression Rating Scale (MADRS) (52), Positive and Negative Syndrome Scale (53), Young Mania Rating Scale (YMRS) (54), Schizo-Bipolar Scale (SBS) (55) and Hollingshead Two-Factor Socioeconomic Rating Scale (SES) (56). Data on average daily chlorpromazine equivalents (avg-daily-cpz) were also recorded for few psychosis participants (49% of 1127 psychosis participants).

Table 1 outlines demographic, cognitive, and clinical characteristics for the B-SNIP study, along with details on analysis datasets (refer Section.2.5) - discovery and replication sets used for hypothesizing and validation.

**Table 1.**
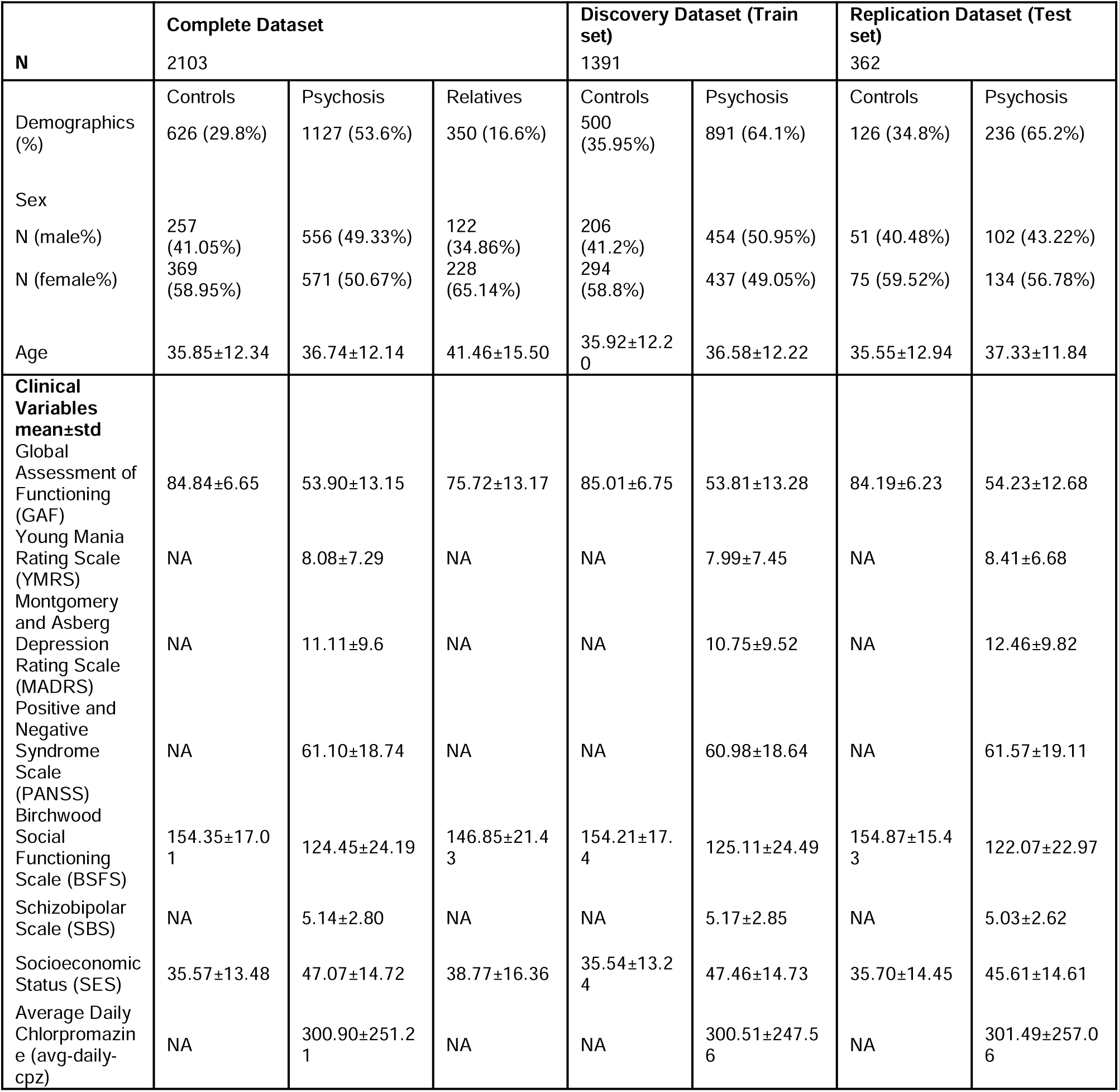
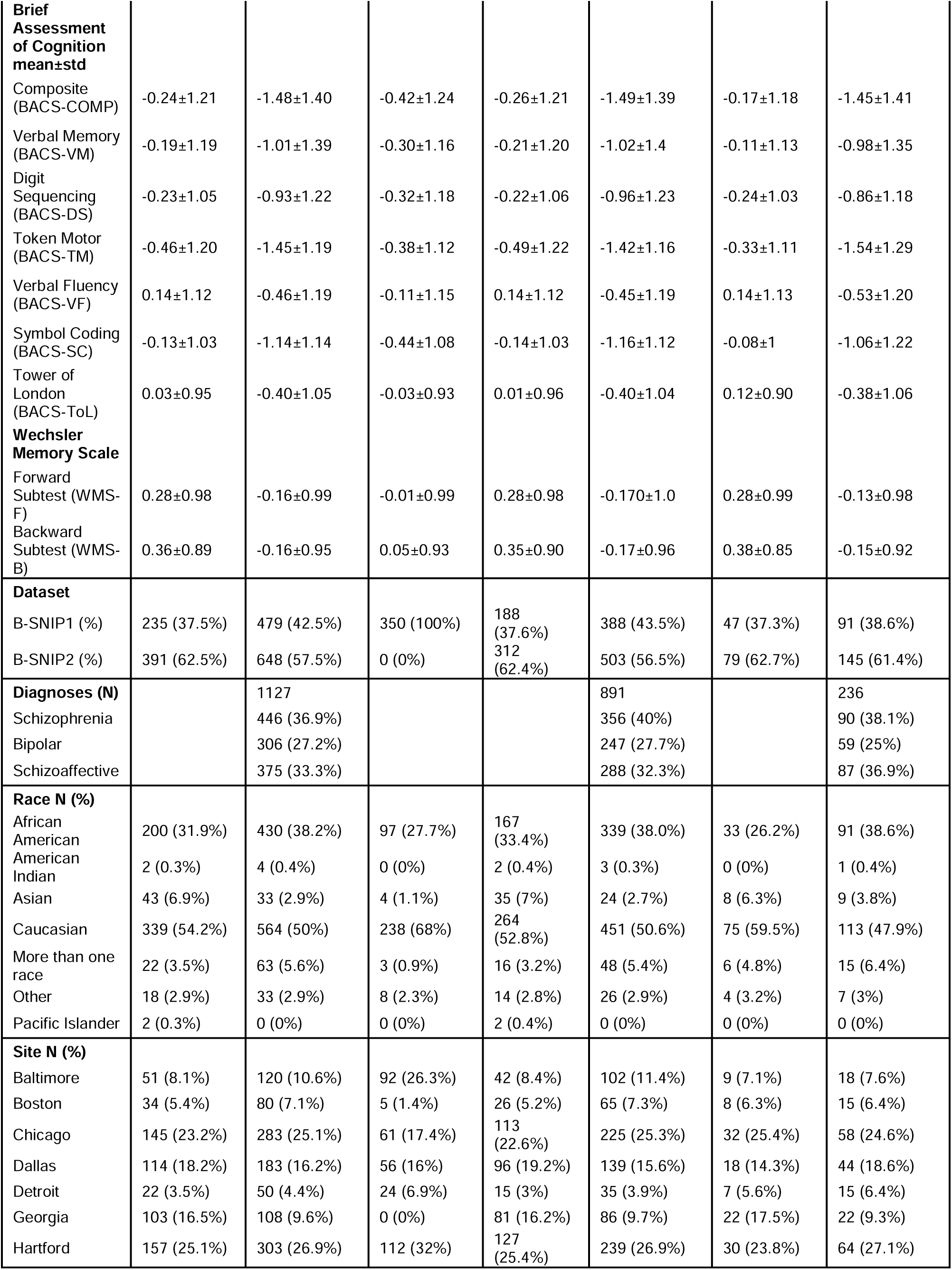
Demographic, clinical, cognitive, diagnostic, and site characteristics of B-SNIP study.

### 2.2. Image Acquisition and Preprocessing

rsfMRI scans were collected from participants across multiple sites using 3T MRI scanners. Preprocessing was conducted using FMRIB Software Library (FSL v6.0; https://fsl.fmrib.ox.ac.uk/fsl/fslwiki) and Statistical Parametric Mapping toolbox (SPM 12; http://www.fil.ion.ucl.ac.uk/spm/) in MATLAB. Preprocessing pipeline corrects for body motion using FSL’s mcflirt function. Distortion correction was performed with FSL’s applytopup function, followed by slice timing correction using SPM toolbox. Scans were warped to Montreal Neurological Institute (MNI) space using an echo planar imaging (EPI) template (57) and resampled to imaging data of 3 × 3 × 3 mm^3^ voxel space through SPM normalization tool. Finally, spatially smoothing was applied using a Gaussian kernel with a full-width half maximum of 6 mm to reduce noise and enhance the signal-to-noise ratio. Refer to Supplementary Section 2 for details on data acquisition parameters and preprocessing.

### 2.3. Estimating subject-specific multi-scale Intrinsic Connectivity Networks (ICNs)

We utilized Group ICA of fMRI Toolbox (GIFT) v4.0c package (https://trendscenter.org/software/gift/) (58) to perform MOO-ICAR (37) and estimate subject-specific multi-scale ICNs. Within this spatially constrained ICA process, we utilized NeuroMark 2.2 template which includes 105 highly reliable ICNs across multiple spatial scales from over 100k+ individuals (38) (Supplementary Section 3; available at https://trendscenter.org/data). These 105 ICNs were grouped and labeled into 7 domains (40) (Fig.1) Cerebellar Domain (CB); Visual Domain (VI) - comprising Occipitotemporal (OT) and Occipital (OC) subdomains; Paralimbic Domain (PL); Subcortical Domain (SC) - Extended Hippocampal (EH), Extended Thalamic (ET), and Basal Ganglia (BG) subdomains; Sensorimotor Domain (SM); Higher Cognition Domain (HC) - Insular Temporal Subdomain (IT), Temporoparietal Subdomain (TP), and Frontal Subdomain (FR); Triple Network Domain (TN) - Central Executive (CE), Default Mode (DM), and Salience (SA) subdomains.

**Fig. 1:**
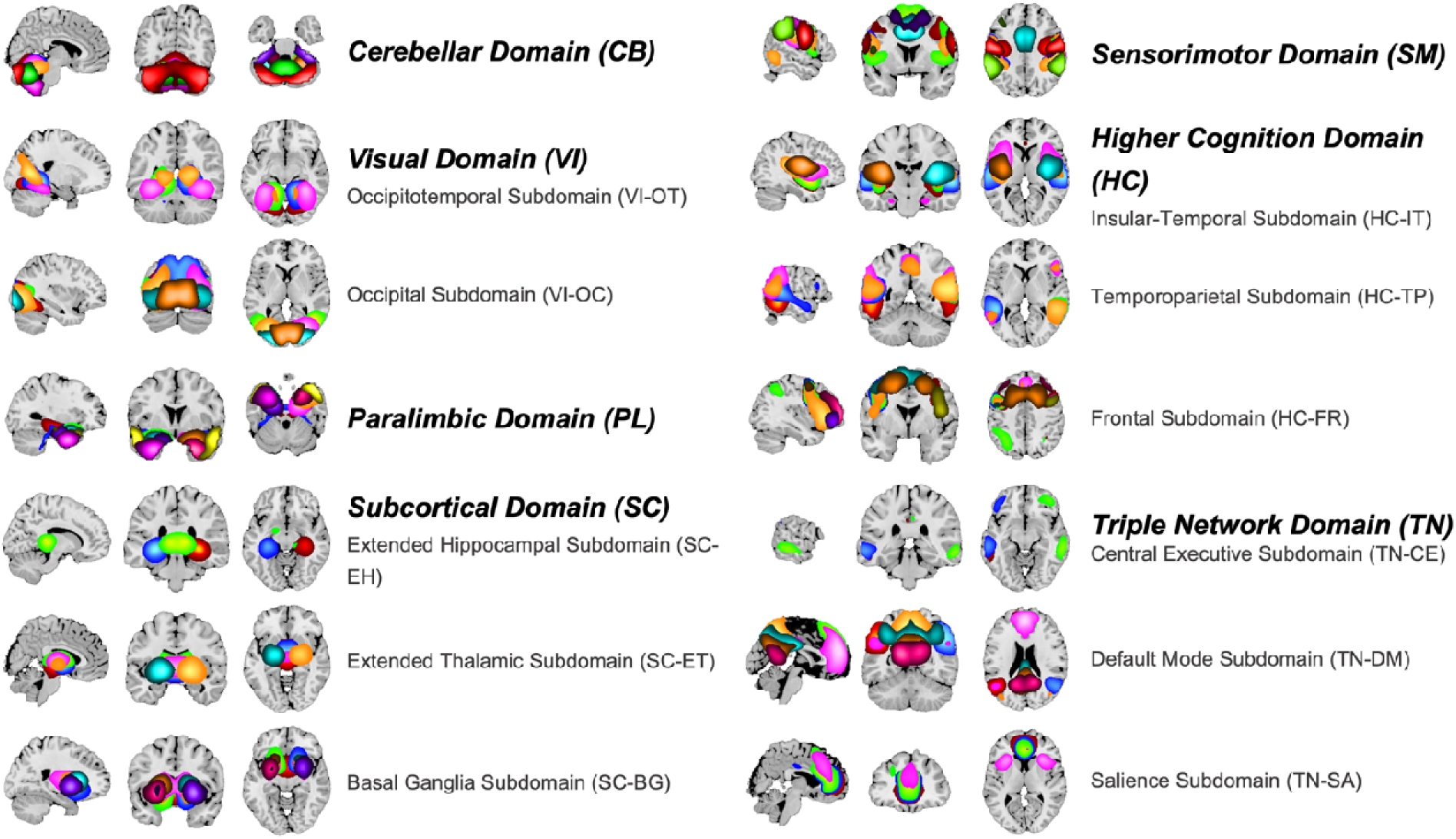
105 Multi-scale Intrinsic Connectivity Networks (ICN) of NeuroMark 2.2 reference template. Figur displays 105 ICN categorized into 7 domains: Cerebellar Domain, Visual Domain (comprising occipitotemporal and occipital subdomains), Paralimbic Domain, Subcortical Domain (comprising Extended Hippocampal, Extended Thalamic and Basal Ganglia subdomains), Sensorimotor Domain, Higher Cognition Domain (comprising Insular Temporal, Temporoparietal and Frontal subdomains) and Triple Network Domain (comprising Central Executive, Default Mode and Salience subdomains). Detailed information on each of the 105 networks can be found i Supplementary Section 3 (40).

### 2.4. Calculating Subject-Specific Multi-Scale Functional Network Connectivity (msFNC)

msFNC calculation involved preprocessing and cleaning the extracted ICN time courses (details in Supplementary Section 4). The whole brain msFNC was estimated from 105 clean time courses by calculating a Pearson correlation between them. Result was a 105 × 105 symmetric msFNC matrix for each participant, with 5460 unique features representing the whole brain multiscale functional connectome (Fig.2A).

**Fig. 2:**
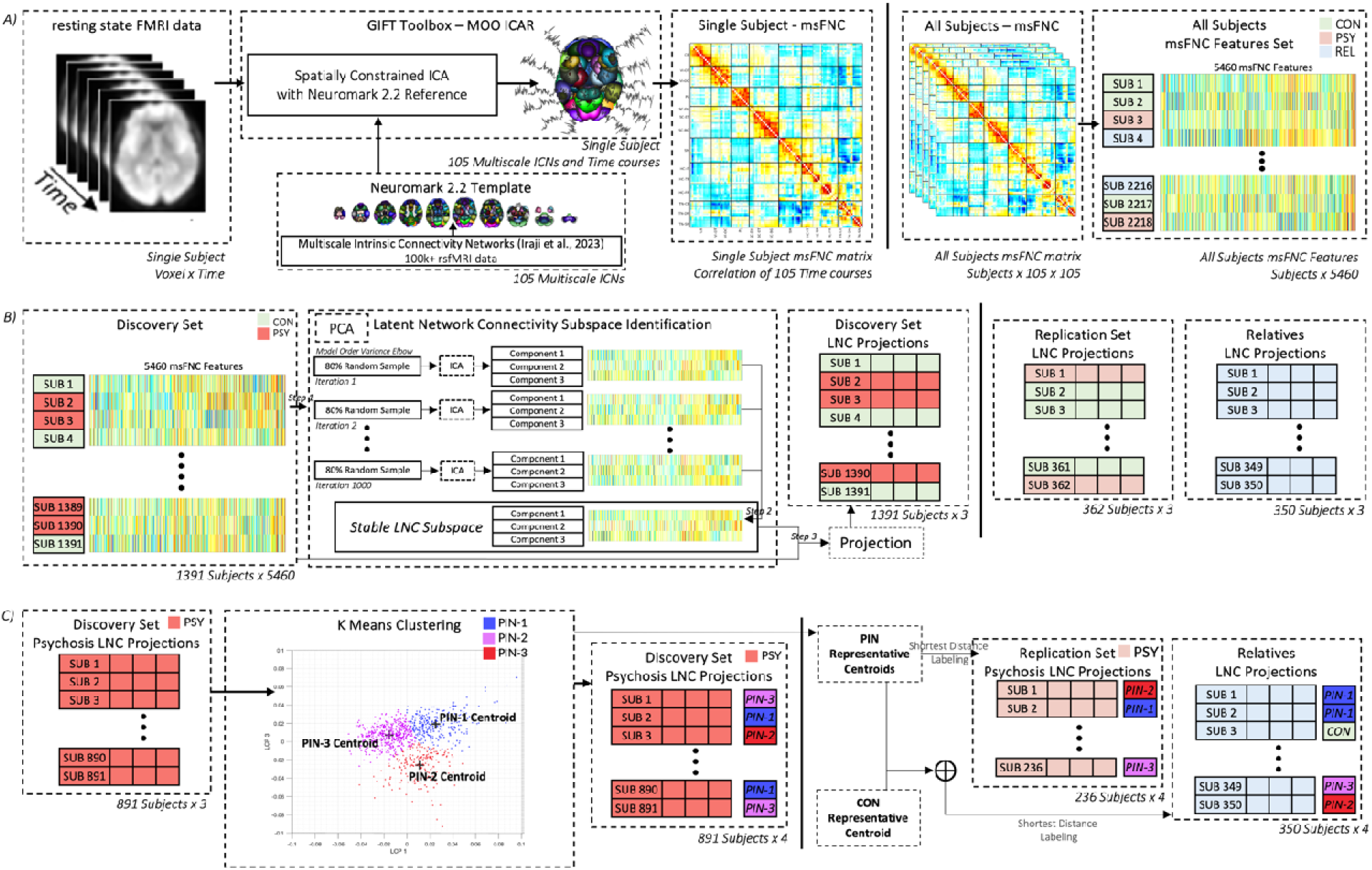
Pipelines illustrating estimation of subject specific multi-scale Functional Network Connectivity (msFNC), extraction of Low-Dimensional Latent Network Connectivity Subspace and identifying of Psychosis Imaging Neurosubtypes (PINs). A) Resting state Functional Magnetic Resonance Imaging (rsfMRI) data for individual subjects is processed using spatially constrained Independent Component Analysis (ICA) with Multivariate Objective Optimization, guided by the NeuroMark 2.2 reference template. This resulted in 105 multiscale Intrinsic Connectivity Networks and their time courses, used to compute a 105 × 105 msFNC matrix. The same procedure is applied across all participants and resulting msFNC matrices are vectorized to extract msFNC features. B) Discovery set contains 80% of entire msFNC data, comprising participants with psychosis and controls. Low-dimensional latent network connectivity (LNC) subspace is extracted by performing ICA on 1000 subsamples of discovery set msFNC. Stable LNC subspace is then estimated from all runs. Then the msFNC of all datasets are projected onto this stable subspace to extract LNC projections. C) LNC projections from participants with psychosis in discovery set are clustered through K-means to identify PINs. Neurosubtypes representative centroids were used to classify participants with psychosis of replication set and assign first-degree relatives to either psychosis neurosubtypes or the control group.

### 2.5. Analysis Datasets: Discovery and Replication Data

We used data from participants with psychosis, their first-degree relatives and controls (data characteristics in Table 1 and Supplementary Section 5). This dataset was randomly divided into discovery and replication sets, comprising 80% and 20% of each group excluding relatives. Both datasets maintained consistent proportion of participants with psychosis and controls. Due to limited avg-daily-cpz information among psychosis participants, the division also ensured comparable sample sizes across both these datasets.

### 2.6. Identification and Characterization of Psychosis Imaging Neurosubtypes

Subsequent analyses focused on extraction of a low-dimensional msFNC subspace using psychosis and control participants, followed by the identification of neurosubtypes based on psychosis participants subspace projections. To prevent systematic biases, we exclusively used discovery dataset for these analyses, while hold-out replication dataset served for validation of the findings.

#### 2.6.1 Extraction of a Low-Dimensional Latent Network Connectivity Subspace

We used a scree plot to identify the dimensionality of subspace that explained the maximum variance in msFNC within discovery set. Subsequently, we employed Infomax ICA (59), which utilizes higher-order statistics to extract independent components and create a low-dimensional latent network connectivity subspace (LNC; Supplementary Section 6). To enhance the reliability and stability of the results, and given the stochastic nature of ICA decomposition, we ran ICA on 1000 subsets of the discovery set using ICASSO technique (60), each containing 80% randomly selected samples. We then selected the most stable LNC components (Fig.2B) across all runs. Finally, high dimensional msFNC data from both discovery and replication datasets are mapped onto the stable LNC subspace. This process reduced data’s dimensionality, resulting in low dimensional latent network connectivity projections (LNC projections).

#### 2.6.2. Identification of Psychosis Imaging Neurosubtypes

We next employed K-means clustering to group the participants with psychosis in discovery set within the LNC subspace (Fig.2C). This clustering used squared Euclidean distance to identify clusters (clustering parameters in Supplementary Section 7). To determine the optimal number of clusters, we applied several techniques, including Calinski-Harabasz index, silhouette, and elbow criteria, all of which indicated that three clusters were optimal. These clusters were designated as PINs, with their centroids serving as representative LNC projections for each neurosubtype group. Finally, new participants from hold-out replication set were assigned to the neurosubtypes based on the minimum squared Euclidean distance between their LNC projections and the neurosubtypes centroids.

#### 2.6.3 Psychosis Imaging Neurosubtypes Cognitive, Clinical Variables, and msFNC Characterization

PINs were characterized using both non-imaging measures, such as cognitive assessments and clinical variables, and imaging measures such as LNC projections and msFNC features. Statistical comparisons were focused on controls versus neurosubtypes and between the neurosubtypes themselves. Linear regression models were used to identify differences between the groups of interest, while controlling for potential confounders, including race, site, mean framewise displacement (mean FD; excluded for cognitive and clinical variable analyses), age, and sex. All analyses were performed on discovery set, with results corrected for multiple comparisons using False Discovery Rate (FDR) correction and validated in hold-out replication set.

#### 2.6.4 Observing Psychosis Participants’ First-Degree Relative’s Intermediate Neurobiology

Considering the shared biology, first-degree relatives of participants with psychosis are expected to exhibit neurobiological patterns intermediate between those of controls and their affected relatives (6,62). We assessed whether the identified PINs could detect these intermediate patterns and compared the results with DSM diagnostic categories.

In this analysis, a representative centroid for the control group was first computed using their average LNC projections from discovery set. Relatives were then assigned to either PINs or the control group based on the minimum squared Euclidean distance to the representative centroids. Same procedure was repeated using DSM diagnoses (SZ, BP, SAD). The results from both analyses (using neurosubtypes and DSM categories) were compared against the null hypothesis and with each other to assess statistical significance.

## RESULTS

Analysis of demographic data revealed no statistically significant age differences between participants with psychosis and controls (Fig.3A; Supplementary Section 5). Distributions of participants, sex (female/male percentages) and DSM categories within psychosis for discovery and replication sets are visualized in pie charts (Fig.3B&C).

**Fig. 3:**
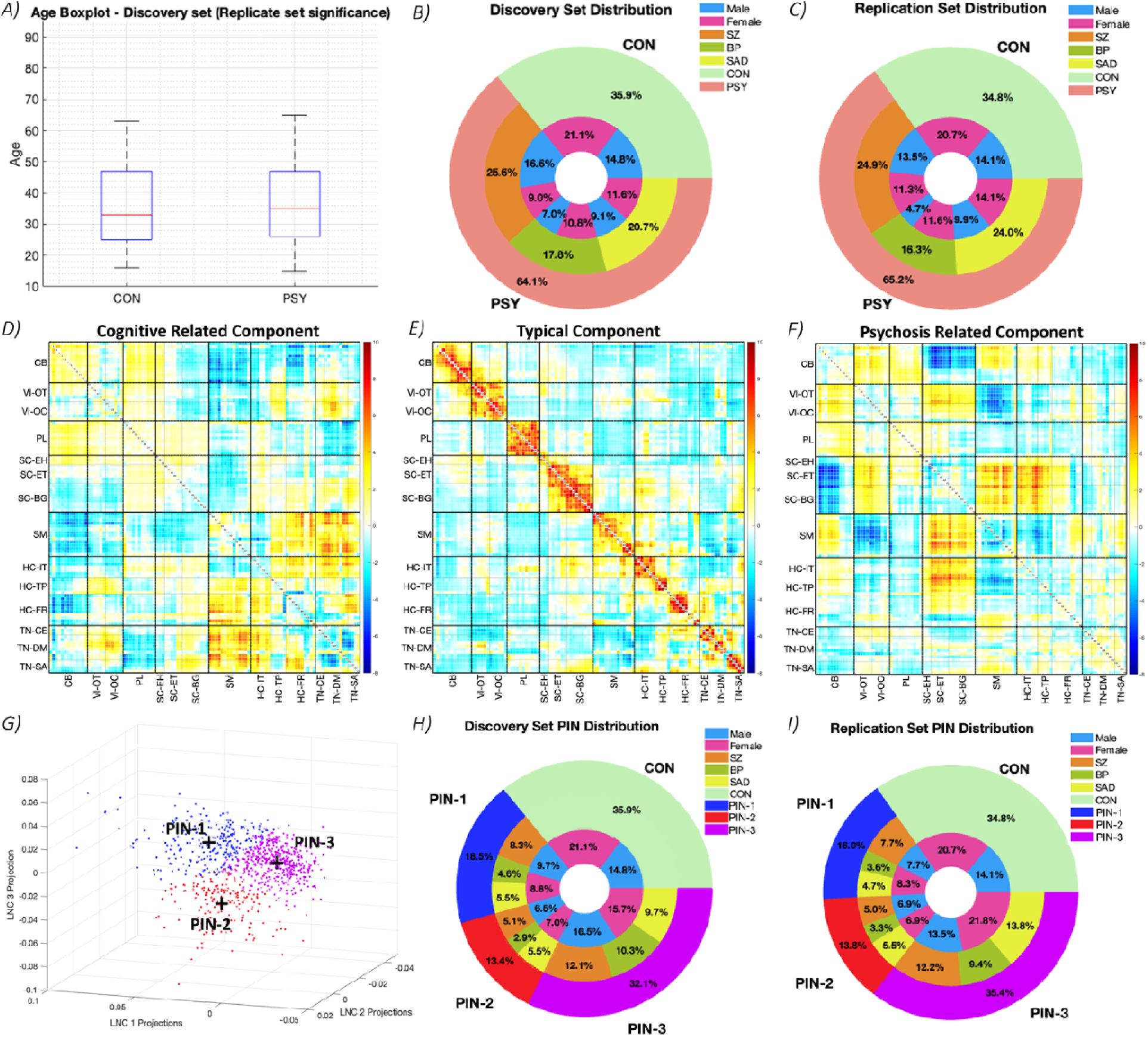
BSNIP Data Distribution, multi-scale Latent Network Connectivity subspace (LNC) and Psychosis Imaging Neurosubtypes (PINs). A) Boxplots represent the age distributions of Healthy Controls (CON) and PSY participants in discovery set. B,C) Pie chart showing BSNIP data distribution in discovery and replication set. The outer pie shows CON, PSY groups distribution, DSM diagnosis distribution indicated in the middle pie, and sex distribution of DSM diagnosis represented within the inner pie. D-F) Subspace of three stable LNC components, each represented by a 105 × 105 matrix. D) Cognitive-Related Component E) Typical Component F) Psychosis-Related Component. Refer Fig.1 and Supplementary Section 3 for additional information on the 105 multi-scale intrinsic connectivity networks. G) PINs LNC projections of all 3 components and centroids are indicated in a 3D scatter plot. This plot indicates PIN-1, PIN-2, and PIN-3. H,I) Pie chart showing BSNIP data distribution in discovery and replication set after identifying PINs. The outer pie shows CON, neurosubtypes distribution, DSM diagnosis along each neurosubtype represented in the middle pie, and the sex distribution of each neurosubtype represented within the inner pie. List of abbreviations – CON: Controls; PSY: Psychosis; LNC: Latent Network Connectivity; SZ: Schizophrenia Disorder; BP: Bipolar Disorder; SAD: Schizoaffective Disorder; PIN: Psychosis Imaging Neurosubtype; PIN-1: Psychosis Imaging Neurosubtype-1; PIN-2: Psychosis Imaging Neurosubtype-2; PIN-3: Psychosis Imagin Neurosubtype-3.

### 3.1. Low-Dimensional Latent Network Connectivity Subspace

Three distinct LNC subspace components were identified (Fig.3D-F): a cognitive-related component (associated with BACS and WMS scores), a typical component (resembling average msFNC patterns), and a psychosis-related component (capturing group differences between controls and psychosis; details in Supplementary Section 6). The extracted LNC components within each run demonstrated high reliability, with ICASSO quality index values (60) of 0.97, 0.97, and 0.96. Additionally, LNC components demonstrated strong consistency across runs, with correlation values exceeding 0.99.

### 3.2 Psychosis Imaging Neurosubtypes

Three-cluster solution was optimal (Supplementary Section 7). Using K-means on LNC projections of psychosis-affected participants, three distinct PINs were identified: PIN-1, PIN-2, and PIN-3 (Fig.3G). Centroids of these neurosubtypes serve as representative values for each group. The distributions of neurosubtypes and their DSM groups are shown for discovery and replication sets (Fig.3H&I).

### 3.3. Psychosis Imaging Neurosubtypes – Non-Imaging Measures Characterization and Validation

#### 3.3.1. Psychosis Imaging Neurosubtypes Cognitive Characterization

Statistically significant differences in all cognitive measures were found between controls and all PINs, with replicable results (Fig.4A; Supplementary Section 8.1). Exceptions included BACS-Tower of London (BACS-ToL) and WMS-Forward subtest scores, where no significant differences were found between controls and PIN-2, and controls and PIN-3 respectively. PIN-1 consistently displayed lower scores compared to PIN-2, which had higher scores, with statistically significant and replicable differences across several cognitive measures including BACS composite score (BACS-COMP), verbal memory, and digit sequencing (Fig.4B). Furthermore, PIN-1 also showed statistically significant, lower scores than PIN-3, particularly in BACS-COMP and BACS-ToL assessment.

**Fig. 4:**
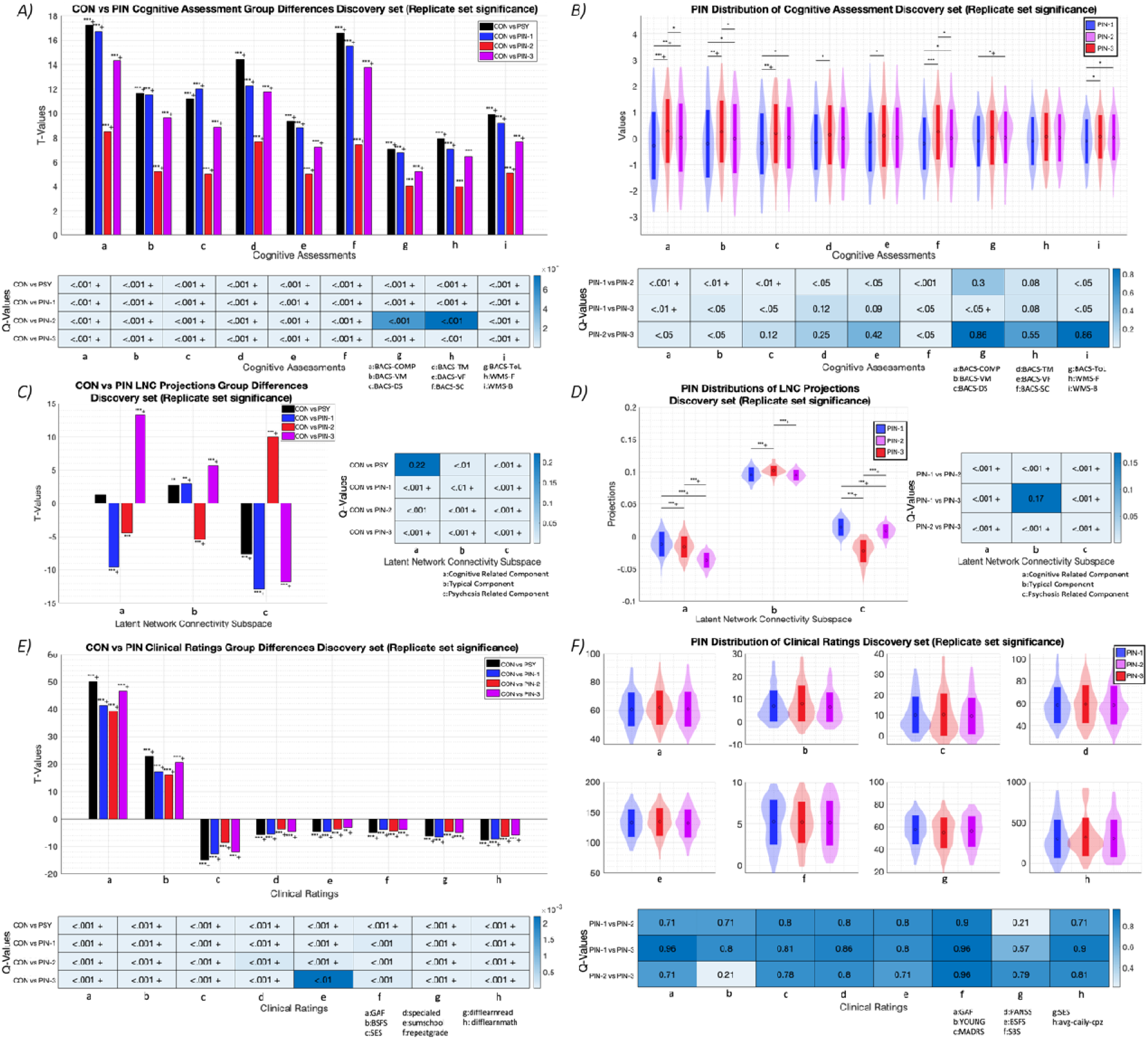
Group Differences between the Control (CON) - Psychosis Imaging Neurosubtypes (PINs) and within PINs across Cognitive Assessment Scores (Brief Assessment of Cognition in Schizophrenia, BACS and Weschler Memory Scale, WMS), Latent Network Connectivity (LNC) Projections and Clinical Variables, in Discovery set. A,B) Different cognitive assessment scores involve BACS Composite Score (BACS-COMP), Verbal Memory (BACS-VM), Token Motor (BACS-TM), Verbal Fluency (BACS-VF), Symbol Coding (BACS-SC), Tower of London (BACS-ToL), WMS Forward (WMS-F) and WMS Backward (WMS-B) subtests. A) Bar plots indicate cognitive assessment scores group differences in discovery set, the x-axis representing different cognitive assessments, and the y-axis showing the T-Values (t-value ranges from 0 to 18) of the group difference. The heatmap below it indicates Q-Values (FDR corrected P-Values) for the group differences in cognitive assessment scores, between CON an PINs in discovery set. B) Violin plots represent PINs distributions of cognitive assessment scores in discovery set, with error bars indicating the mean and standard deviation. The heatmap below indicates the Q-Values for group differences in cognitive assessment scores, within neurosubtypes in discovery set. C,D) Different LNC projections involve Cognitive-Related Component projections (LNC projections 1), Typical Component projections (LNC projections 2), and Psychosis-Related Component projections (LNC projections 3). C) Bar plots indicate different LNC projections group differences in discovery set, with the x-axis representing different LNC component projections respectively, and the y-axis showing the T-Values (t-value ranging from-15 to 15) of the group differences. The heatmap adjacent to it indicates the Q-Values (FDR corrected P-Values) of the group differences in LNC projections, between CON and PINs of discovery set. D) Violin plots represent the PINs distributions of LNC projections in discovery set, with error bars indicating the mean and standard deviation. The heatmap adjacent to it indicates the Q-Values of the group difference in LNC projections, within neurosubtypes of discovery set. E,F) Different clinical variables involve Global Assessment of Functioning (GAF), Young Mania Rating Scale (YMRS), Montgomery-Asberg Depression Rating Scale (MADRS), Positive and Negative Syndrome Scale (PANSS), Birchwood Social Functioning (BSFS), Schizo-Bipolar Scale (SBS), Socioeconomic Rating Scale (SES), and Average Daily Chlorpromazine Equivalents (avg-daily-cpz). E) Bar plots indicate group differences across clinical variables in discovery set, with the x-axis representing different clinical variables and the y-axis showing the T-Values (t-value ranging from-20 to 60) of the group differences. The heatmap below it indicates Q-Values (FDR corrected P-Values) of the group differences in clinical variables, between CON and PINs of discovery set. F) Violin plots represent PINs distributions of clinical variables in discovery set, with error bars indicating the mean and standard deviation. The heatmap below it indicates Q-Values of the group difference in clinical variables, within neurosubtypes of discovery set. A,C,E) Plots indicating group differences between CON and PINs, using color bars to show different group comparisons (black = CON vs. PSY group, blue = CON vs. PIN-1, red = CON vs. PIN-2, and violet = CON vs. PIN-3). B,D,F) Plots indicating distributions within PINs uses blue, red, and violet to indicate PIN-1, PIN-2, and PIN-3 distributions respectively. A-F) In all the plots, Single * represents statistical significance and q-value < 0.05, Double ** represents q-value < 0.01, Triple *** represents q-value < 0.001, and + indicates the test significance in replication set. All heatmaps indicating + represent test significance in replication set. List of abbreviations – CON: Controls; PSY: Psychosis; LNC: Latent Network Connectivity; PIN: Psychosis Imaging Neurosubtype; PIN-1: Psychosis Imaging Neurosubtype-1; PIN-2: Psychosis Imaging Neurosubtype-2; PIN-3: Psychosis Imaging Neurosubtype-3; BACS: Brief Assessment of Cognition; BACS-COMP: Composite Score; BACS-VM: Verbal Memory; BACS-DS: Digit Sequencing; BACS-TM: Token Motor; BACS-VF: Verbal Fluency; BACS-ToL: Tower of London; WMS-F: Wechsler Memory Scale; WMS-F: Forward Subtest; WMS-B: Backward Subtest; GAF: Global Assessment of Function; YMRS: Young Mania Rating Scale; MADRS: Montgomery-Asberg Depression Rating Scale; PANSS: Positive and Negative Syndrome Scale; BSFS: Birchwood Social Functioning; SBS: Schizo-Bipolar Scale; SES: Socioeconomic Rating Scale; avg-daily-cpz: Average Daily Chlorpromazine Equivalents

#### 3.3.2 Psychosis Imaging Neurosubtypes Clinical Characterization

In the discovery set, PINs were significantly associated with DSM diagnoses (X^2^=13.9, p-value<0.001), but this relationship was not replicated in the replication set (X^2^=3.6, p-value > 0.1). Although DSM diagnoses were unevenly distributed across neurosubtypes, all diagnoses were represented in each group (Fig.3B& C; Table 2). PIN-1 and PIN-3 were predominantly composed of SZ participants, while PIN-2 had highest proportion of SAD participants.

**Table 2.**
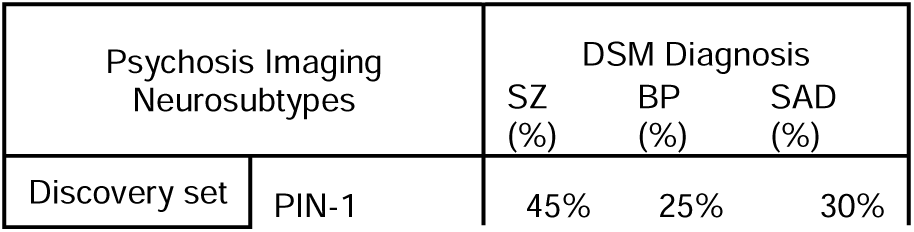
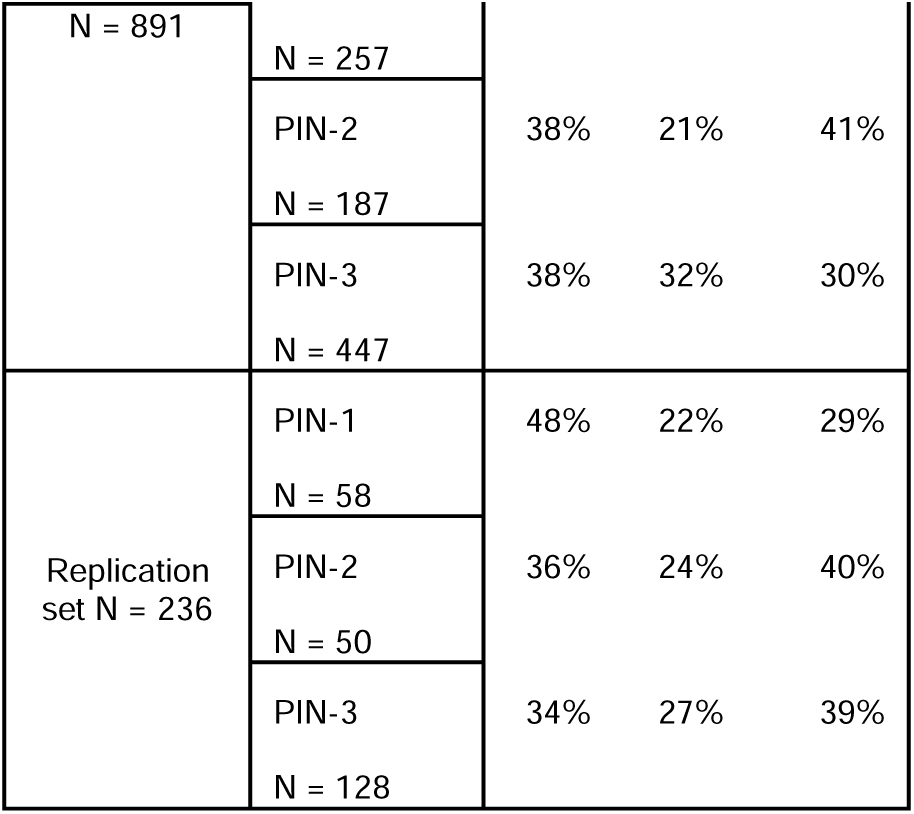
Relation between Psychosis Imaging Neurosubtypes and DSM Diagnosis.

Clinical variables for controls showed statistically significant differences compared to PINs with replicable results (Fig.4E). No statistically significant differences were observed between neurosubtypes in these variables (Fig.4F; Supplementary Section 8.3), after correcting for multiple comparisons.

### 3.4. Psychosis Imaging Neurosubtypes – Imaging Measures Characterization and Validation

#### 3.4.1. Psychosis Imaging Neurosubtypes LNC Projections Characterization

Significant and replicable group differences were observed between controls and all PINs across the projections of cognitive-related, typical, and psychosis-related components (Fig.4C; Supplementary Section 8.2). However, controls and PIN-2 were not significantly different for the cognitive-related component. Statistically significant differences were also found between neurosubtypes with replicable results, except between PIN-1 and PIN-3, which showed no significant difference along the typical component projections (Fig.4D).

#### 3.4.2. Psychosis Imaging Neurosubtypes msFNC Characterization

Statistical comparisons of msFNC between controls and PINs, as well as within psychosis PINs, revealed differences in discovery sets, which were replicated in replication set (summary in Fig.5). Strong correlations of T-values between groups in discovery and replication sets (t-value similarities were rho_(CON,PIN-1)_ = 0.85, rho_(CON,PIN-2)_ = 0.66, rho_(CON,PIN-3)_ = 0.84, rho_(PIN-1,PIN-2)_ = 0.87, rho_(PIN-1,PIN-3)_ = 0.87, rho_(PIN-2,PIN-3)_ = 0.90) further confirms the replicability of the findings. Summary below highlights a few of the differences, including both within and between domain connectivity variations, with more details provided in Supplementary Section 8.4.

**Fig. 5.**
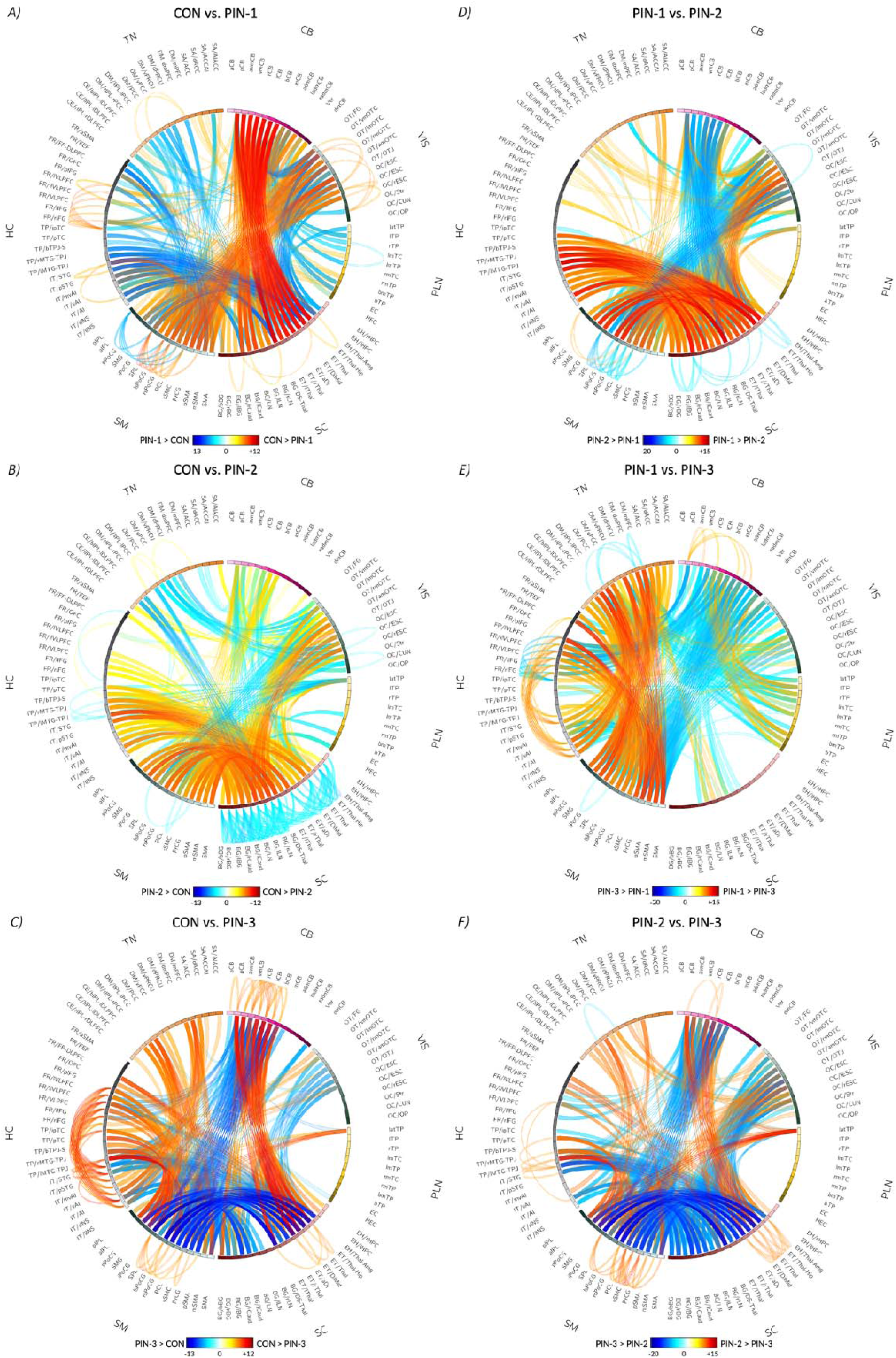
: **Group Differences between Controls - Psychosis Imaging Neurosubtypes (PINs), and within PINs along Multi-scale Functional Network Connectivity (msFNC): Replicable Differences of Discovery set.** msFNC represents connectivity between 105 Multi-scale Intrinsic Connectivity Networks (msICNs). Refer Fig.1 and Supplementary Section 3 for additional information on the 105 multi-scale intrinsic connectivity networks. Connectograms illustrate the top 10% of statistically significant connectivity differences in discovery set, highlighting only those that were replicable within replication set. Complete msFNC differences can be found in Supplementary Section 8.4. Outward links represent within domain connectivity differences, while inward links indicate between domain differences, with link intensity reflecting the T-values. A-C) Connectogram showing common group differences, between CON vs. PIN-1, CON vs. PIN-2, CON vs. PIN-3 respectively. All links indicate t-value of difference, having same range of-13 to 12. D-F) Connectogram showing common group differences, between PIN-1 vs. PIN-2, PIN-1 vs. PIN-3, and PIN-2 vs. PIN-3 respectively. All links indicate t-value of difference, having same range of-21 to 16. List of abbreviations – CON: Controls; PSY: Psychosis; LNC: Latent Network Connectivity; PIN: Psychosis Imaging Neurosubtype; PIN-1: Psychosis Imaging Neurosubtype-1; PIN-2: Psychosis Imaging Neurosubtype-2; PIN-3: Psychosis Imaging Neurosubtype-3.

##### 3.4.2.1. Psychosis Imaging Neurosubtypes - msFNC Patterns within Functional Domains

**Cerebellar Domain:** PIN-3 showed hypoconnectivity in comparison with controls across all ICNs within CB domain (Fig.5C), and no significant differences between PIN-1, PIN-2 compared with controls (Fig.5A&B). PIN-1 exhibited increased connectivity compared to PIN-3, specifically between ICN 2 (posterior cerebellum) and entire CB domain (Fig.5E). PIN-2 also displayed higher connectivity compared to PIN-3 across the entire domain (Fig.5F).

**Subcortical Domain:** PIN-1 and PIN-3 showed hypoconnectivity in comparison with controls, across BG subdomain and EH, ET subdomains respectively (Fig.5A-C). However, PIN-2 exhibited hyperconnectivity compared to controls, particularly in connectivity between ICN 39 and ICN 40 (overlapping with thalamus, hippocampus, and amygdala networks) as well as ICNs 50 and 54 (overlapping with left caudate and bilateral basal ganglia networks; Fig.5B). PIN-1 showed reduced connectivity compared to PIN-2 (Fig.5D) and PIN-2 exhibited increased connectivity when compared with PIN-3 (Fig.5F) across SC domain.

**Higher Cognition Domain:** Within Higher Cognition (HC) domain, PIN-1 displayed hypoconnectivity in comparison to controls within Insular Temporal (IT) subdomain, particularly between ICN 80 (inferior posterior temporal cortex) from this subdomain and ICNs from Frontal (FR) subdomain (Fig.5A). PIN-3 also exhibited hypoconnectivity compared to controls, between IT and FR subdomains, specifically between ICN 76 (left middle temporal gyrus, temporoparietal junction) and ICN 80 of TP subdomain with ICNs from IT subdomain (Fig.5C). Psychosis PIN-1 and PIN-2 did not show any significant connectivity differences within HC domain (Fig.5D). Both PIN-1 and PIN-2 exhibited increased connectivity compared to PIN-3 (Fig.5E&F) in IT, FR subdomain connectivity. But PIN-1 displayed lower connectivity compared to PIN-3 between ICN 80 and FR subdomain (Fig.5E).

##### 3.4.2.2. Psychosis Imaging Neurosubtypes - msFNC Patterns between Functional Domains

**Cerebellar - Subcortical Domains Connectivity:** PIN-1 and PIN-3 demonstrated an overall hypoconnectivity in comparison to controls (Fig.5A-C), but PIN-2 hyperconnectivity (Fig.5B) was observed when compared to controls, particularly in ICN 2 (posterior cerebellum) and ICN 3 (anterior ventromedial cerebellum) connected with SC - ET and BG subdomains. PIN-2 displayed increased connectivity than PIN-3 (Fig.5F), while PIN-1 exhibited reduced connectivity with respect to PIN-2 (Fig.5D).

**Cerebellar - Sensorimotor Domains Connectivity:** PIN-3 showed hyperconnectivity compared to controls, but PIN-1 and PIN-2 displayed hypoconnectivity in comparison to controls (Fig.5A-C) within CB domain ICNs 9 - 13 and SM domain (dorsomedial cerebellum networks). PIN-2 displayed lower connectivity compared with PIN-3, and PIN-1 exhibited lower connectivity compared to PIN-3 (Fig.5E-F). However, PIN-1 and PIN-2 reveal minimal connectivity differences, except for PIN-1 exhibiting higher connectivity in ICN 2,5,6 (posterior, left, right cerebellum networks) connectivity with SM (Fig.5D).

**Visual - Subcortical Domains Connectivity:** PIN-1 and PIN-3 showed hyperconnectivity compared to controls (Fig.5A&B), but PIN-2 showed hypoconnectivity against controls, particularly in ET subdomain (ICNs 41 - thalamus, 44 - right thalamus) and BG subdomain (ICNs 46 - 49 involving dorsal striatum and lentiform networks). PIN-1 exhibited increased connectivity in contrast to PIN-2 (Fig.5D), while PIN-2 exhibited reduced connectivity compared to PIN-3 (Fig.5F).

**Visual - Sensorimotor Domains Connectivity:** PIN-1 and PIN-3 exhibited hypoconnectivity against controls (Fig.5A&C), involving VI domain connectivity with ICNs 59 - 64 of SM domain (postcentral gyrus, superior sensorimotor cortex, paracentral lobule; Fig.5C). However, PIN-2 showed hyperconnectivity compared to controls (ICNs 59 - 63; Fig.5B). PIN-1 showed reduced connectivity compared to PIN-2 and PIN-3 (Fig.5D&E), while PIN-2 displayed increased connectivity with respect to PIN-3 (Fig.5F).

**Subcortical - Sensorimotor Domains Connectivity:** PIN-1 and PIN-3 exhibited hyperconnectivity against controls (Fig.5A&C). In contrast, PIN-2 displayed hypoconnectivity in comparison to controls, particularly in SC - ET, BG subdomain connectivity with SM (Fig.5B). Among neurosubtypes, PIN-1 and PIN-3 exhibited dysconnectivity (Fig.5E). However, PIN-1 exhibited higher connectivity than PIN-2 (Fig.5D), while PIN-2 displayed lower connectivity in comparison to PIN-3 (Fig.5F).

**Higher Cognition - Subcortical Domains Connectivity:** PIN-2 displayed hypoconnectivity with respect to controls in HC connectivity with SC - ET, BG subdomains (Fig.5B). However, both PIN-1 and PIN-3 exhibited hyperconnectivity compared to controls between HC - IT, TP and SC, while showing hypoconnectivity in HC - FR and SC connectivity (Fig.5A&B). PIN-1 showed higher connectivity when compared to PIN-2 in HC and SC - ET, BG connectivity (Fig.5D). PIN-2 exhibited lower connectivity compared to PIN-3 in HC - IT, TP connectivity with SC, but displayed increased connectivity in comparison with PIN-3 in HC - FR connectivity with SC (Fig.5F).

### 3.5. Psychosis First-Degree Relatives Intermediate Neurobiology

Out of the 350 relatives, 90.29% were classified into PINs (Fig.6A- C) and 55.06% among those matched neurosubtype of affected family members, showing a statistical significance above the null (p-value_Relatives-to-PIN-1_ < 0.001, p-value_Relatives-to-PIN-2_ < 0.05, p-value_Relatives-to-PIN-3_ < 0.001). To address potential confounding effects of age differences, age was regressed out from msFNC data of relatives. Subsequent classification using adjusted LNC projections still produced significant results with even lower p-values (p-value_Relatives-to-PIN-1_ < 0.001, p-value_Relatives-to-PIN-2_ < 0.01, p-value_Relatives-to-PIN-3_ < 0.001). Details on msFNC differences of relatives based on this classification are in Supplementary Section 9.

**Fig. 6.**
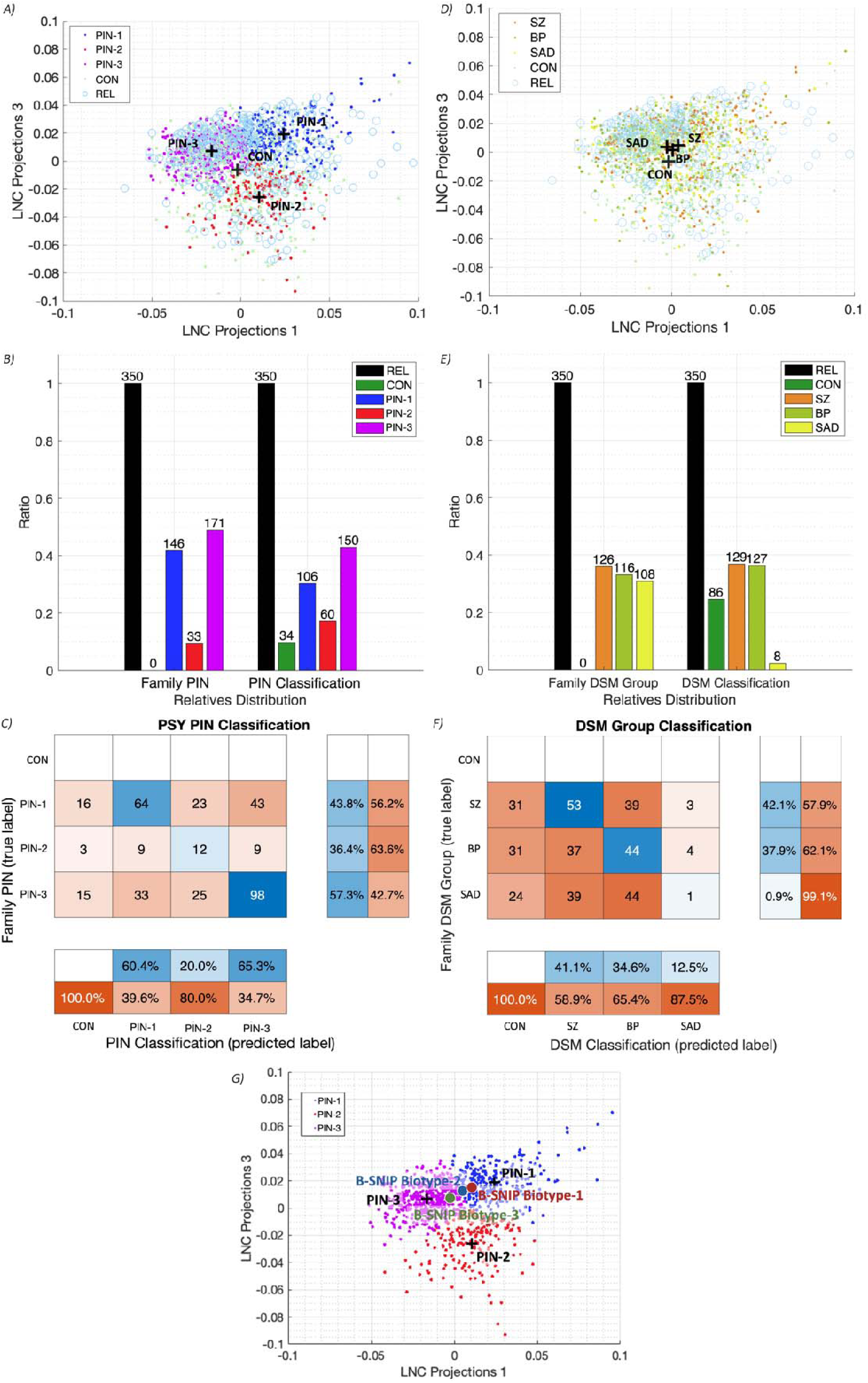
: **Classification of Psychosis first-degree Relatives (REL) based on Psychosis Imaging Neurosubtypes (PINs), DSM groups; Comparison of PINs with B-SNIP Biotypes** A-C) Results of REL participants classification based on PINs and control group. A) Scatter plot showing Latent Network Connectivity (LNC) projections of Healthy Controls (CON), PINs of discovery set, and their first-degree REL participants. Neurosubtypes and CON centroids are also highlighted. B) Bar plot showing distribution of REL participants affected family member’s PIN and CON group/PIN classification. C) Confusion matrix comparing REL participant’s family member’s PIN (true label), with their PIN classified label (predicted label). D-F) Results of REL participants classification based on DSM groups and control groups. D) Scatter plot showing LNC projections of Healthy Controls (CON), DSM groups of discovery set, and their first-degree REL participants. DSM group and CON centroids are also highlighted. E) Bar plot showing distribution of REL participants affected family member’s DSM group and CON/DSM group classification. F) Confusion matrix comparing REL participant’s family member’s DSM group (true label), with their DSM group classified label (predicted label). G) Comparison of PINs (PIN-1, PIN-2 and PIN-3) with B-SNIP study identified Biotypes (Biotype-1, Biotype-2 and Biotype-3) (6). Scatter plot showing Latent Network Connectivity (LNC) projections of PINs and B-SNIP Biotype groups average LNC projections highlighted. List of abbreviations – CON: Controls; PSY: Psychosis; REL: Psychosis participants first-degree relatives; LNC: Latent Network Connectivity; SZ: Schizophrenia Disorder; BP: Bipolar Disorder; SAD: Schizoaffective Disorder; PIN: Psychosis Imaging Neurosubtype; PIN-1: Psychosis Imaging Neurosubtype-1; PIN-2: Psychosis Imaging Neurosubtype-2; PIN-3: Psychosis Imaging Neurosubtype-3.

When relatives were classified based on DSM diagnoses (Fig.6D-F), 75.43% were assigned to DSM categories, but only 37.12% matched the diagnoses of their affected family members (Fig.6F). This classification was not statistically significant (p- value_Relatives-to-SZ_ = 0.1008; p-value_Relatives-to-BP_ = 0.2016; p-value_Relatives-to-SAD_ = 0.9330).

## DISCUSSION

The current study adopts a data-driven approach focusing solely on neurobiological perspectives to identify imaging neurosubtypes within psychosis. By employing robust spatially constrained ICA referencing NeuroMark 2.2, the study identified accurate subject-level brain-wide msFNC estimations, enabling a comprehensive examination of psychosis heterogeneity. The study utilized rsfMRI data to identify distinct PINs, contributing to ongoing efforts to better understand mental disorders (5).

Through extraction of a low-dimensional msFNC subspace, critical latent patterns of functional features were captured (LNC subspace and projections), allowing the identification of PINs. These neurosubtypes were consistent even after accounting for confounding variables (Supplementary Section 7). Importantly, neurosubtypes demonstrated larger msFNC differences compared to controls than when analyzing entire psychosis group or individual DSM based groups (Supplementary Section 10), revealing a reduction in heterogeneity.

Distinct PINs not only exhibited unique connectivity traits but also revealed significant cognitive performance variations across themselves. These findings suggest a link between latent connectivity patterns, rooted within multiscale information, and cognitive performance. This framework may offer a pathway toward targeted, personalized treatments informed by connectivity-based profiles, advancing our understanding of psychosis and enhancing the precision of interventions to address its diverse manifestations and emphasizes the importance of moving beyond traditional symptom-based approaches toward data-driven neurobiological methods that better capture psychosis heterogeneity.

First-degree relatives exhibited intermediate connectivity patterns aligning significantly with their affected family members and showing similar msFNC differences compared to controls. Conversely, DSM-based analysis showed weaker alignment and lacked statistical significance, further emphasizing the effectiveness of neurobiological methods in capturing these relationships.

### 4.1. Psychosis Imaging Neurosubtypes Cognitive Characterization

Each PIN defined by a distinct connectivity profile, revealed significant variations in cognitive performance. PIN-1 was most impaired, followed by PIN-3, while PIN-2 demonstrated better preserved cognitive abilities (Fig.4A; Supplementary Fig.6A). This pattern of cognitive performance— controls > PIN-2 > PIN-3 > PIN-1, was consistent across discovery and hold-out replication sets, strengthening the reliability of the findings. PIN-2 despite showing cognitive deficits, was closer to controls in some of the cognitive measures, highlighting its comparatively preserved cognitive profile. Connectivity analyses supported this trend: PIN-1 and PIN-3 exhibited the greatest connectivity deviations from controls, while PIN-2 showed the least. Specifically, PIN-1 demonstrated pronounced hypoconnectivity in CB-SC, VI-SM domains and hyperconnectivity in VI-SC, VI-CB, and SC-SM domains. Similar patterns were observed in recent cognitive-focused study (16), which identified two cognitive biotypes with distinct connectivity profiles. The worst-performing cognitive biotype in that study aligned with our PIN-1 in connectivity patterns, while its preserved cognitive biotypes shared features with our PIN-3, including hypoconnectivity in HC-SM and HC-SC and hyperconnectivity in CB-SM and SC-SM. Importantly, our approach not only replicated known patterns but also uncovered PIN-2, which exhibited unique connectivity profile, and was a more preserved group. Given the likelihood of multiple subgrouping ways depending on the clinical or research objective (63), these two different approaches, along with previous efforts (6) may offer complementary insights and utility.

### 4.2. Psychosis Imaging Neurosubtypes Clinical Characterization

No significant differences were noted in clinical variables among PINs. PIN-1 presented a higher proportion of SZ participants (45%) and PIN-2 of SAD participants (40%), a pattern consistent in replication. The neurobiological approach effectively grouped participants with homogenous connectivity patterns, revealing this imbalance in DSM diagnoses while highlighting cognition as a key differentiating factor. These findings emphasize the heterogeneity within DSM diagnoses and while these remain important for clinical classification, they may not fully align with underlying neurobiological connectivity profiles.

### 4.3. Psychosis Imaging Neurosubtypes msFNC Characterization

Previous studies reported psychotic individuals presenting hypoconnectivity between CB-SC regions and hyperconnectivity between SC-HC, SC-SM regions (11,39,64,65). In this study, these trends were stronger in both PIN-1 and PIN-3 (Fig.5A& B; Supplementary Section 8.4). Conversely, PIN-2 showed an opposite pattern of these connections, supporting evidence that suggests possible organizations in brain oscillations that are essential for cognitive function (66,67).

Some brain regions that showed connectivity differences between PINs in the HC domain (IT, TP, and FR subdomains) are closely related to auditory and language-related brain functions (68,67,69). These connectivity variations (Fig.5; Supplementary Fig.8) in PIN-2 could be linked to its distinct typical and psychosis-related component projection values findings, as well as its better cognitive performance, suggesting that these differences might reflect compensatory or adaptive mechanisms.

PIN-1 and PIN-3 showed hyperconnectivity between sensory-related domains (i.e., VI-SC as well as VI-SM) which has been implicated as a pattern characteristic of psychosis (70). However, PIN-2 displayed hypoconnectivity between VI-SC domains, with no connectivity differences between VI-SM domains (71). This is consistent with PIN-2’s low psychosis-related component projections and high typical component projections, which may suggest adaptive neurobiological processes that might help mitigate psychosis symptoms or enhance cognitive resilience.

### 4.4. Psychosis First-Degree Relatives Intermediary Neurobiology

First-degree relatives had intermediate connectivity patterns between controls and their affected family members. About 55% of relative participants matched their family’s PIN, which was statistically significant. Connectivity differences in relatives were weaker but similar to those observed between neurosubtypes and controls. These findings align with previous studies (26,72) and emphasize the importance of investigating genetic and familial risk factors in psychosis. In contrast, only 37% of relatives matched the same DSM diagnosis as their family members, which was not statistically significant. This highlights the biological heterogeneity of psychosis and neurosubtypes effectiveness in capturing shared neurobiological traits, which could offer a more precise framework for understanding the disorder (26).

### 4.5. Psychosis Imaging Neurosubtypes Compared with B-SNIP Biotypes

This study utilizes rsfMRI to identify psychosis neurosubtypes based on intrinsic brain function, contrasting with the B-SNIP study (6), which employed a combination of cognitive, eye movement and EEG data. The three B-SNIP Biotypes appear to reflect variations in cognitive deficits, with B-SNIP Biotype-1 and 3 exhibiting the most and least impairment, respectively. This finding is expected as cognitive performance variables were used in clustering. Notably, our PINs, derived purely from intrinsic brain function, also suggest distinct cognitive differences between neurosubtypes, indicating that cognition is a key clinical dimension in psychosis.

To further explore this relationship, we identified the centroids of B-SNIP Biotypes within the LNC subspace. Our analysis revealed that B-SNIP Biotype-1 is closest to our PIN-1, which also demonstrated the most impaired cognitive performance (Fig.6G). Notably, both PIN-1 and B-SNIP Biotype-1 groups in these studies consisted of higher SZ participants. Additionally, the projection of B-SNIP Biotype-3, which exhibited better cognitive performance compared to other Biotypes, was closest to our PIN-3. Overall, B-SNIP Biotypes-1 through 3 were well-positioned within the intrinsic LNC subspace along a continuum defined by our PIN-1 and 3, showing a certain level of consistency in terms of clinical interpretation. All these together, further emphasize the link between brain function and cognition and the key role of cognition in psychosis, underscoring the need for further investigation.

### 4.6. Low-Dimensional Latent Network Connectivity Subspace

PIN-1, the most severely cognitively affected group, exhibited severe psychosis-related component projections (indicative of severe psychosis) and high cognitive-related component projections (reflecting poor cognition). Conversely, PIN-2 exhibited better cognitive performance, with lower projections in psychosis-related and cognitive-related components, and highest typical component projections, indicating strong within-domain connectivity. Low-dimensional subspace of multiscale information provided key characterizations of neurosubtypes and offers promising translational potential. It could help refine psychosis diagnostics by projecting affected individuals’ msFNC to the subspace, which allows uncovering their latent profiles and informing interventions relevant for developing personalized treatment strategies.

## LIMITATIONS AND FUTURE WORK

Our study explored psychosis heterogeneity through neurobiological measures, specifically brain-wide functional connectivity, to assess its ability to define and characterize psychosis neurosubtypes. While connectivity offers valuable insights, structural, genetic and environmental factors also play crucial roles in the etiology and progression of psychosis (73–77). Future research should aim to integrate more psychosis-related factors using multi-modal methodologies (78,79) alongside other and more diverse datasets (80,81) to achieve a more comprehensive understanding of psychosis.

Sex differences can significantly influence psychosis pathophysiology (82). While our analysis corrects for sex, it requires thorough exploration to better understand the impact of sex on the pathophysiology (83,84). Future investigations could also benefit from incorporating dynamic functional connectivity, particularly over multiscale approaches, which have shown promise in sex differences to clinical symptoms and genetic risk in psychosis (39,84–86).

Additional limitations include the lack of evaluations on potential confounders such as medication dosages (antipsychotics, mood stabilizers, antidepressants, and anticholinergics), substance use (cannabis, alcohol and nicotine) and medical comorbidities such as cardiometabolic disturbances, all of which potentially influence brain connectivity and cognition (31,87–90). B-SNIP psychosis participants are symptomatically stable, and clinical characterization revealed no differences in PANSS, MADRS, YMRS, which could be possibly related to treatment improvement. Furthermore, our Caucasian-dominant cohort (60% of participants) limits the generalizability of results to diverse populations.

## CONCLUSION

Our study assessed the potential of intrinsic brain activity in identifying PINs, addressing the heterogeneity of psychosis and advancing the understanding of its spectrum. The neurosubtypes’ replicable connectivity patterns and cognitive profiles highlight the effectiveness of low-dimensional subspace analysis informed by brain-wide multiscale connectivity data. Relatives displayed comparable neurobiology to their affected family members, highlighting genetic and familial contributions to psychosis. Within our findings, cognition emerged as a core clinical dimension of psychotic disorders emphasizing cognitive dysfunction remedy as a relevant unmet need. These neurosubtypes may serve as useful for stratified targeted cognitive treatments for psychosis offering promising avenues for future research and therapeutic innovation.

## ACKNOWLEDGEMENTS

AI has received grant support from the National Institutes of Health (R01MH136665) and Georgia State University’s Research Initiation Grant (RIG) program. VDC has received grant support from the National Institutes of Health (R01MH123610) and National Science Foundation (2112455).

## AUTHOR CONTRIBUTIONS

RB had conducted the study, designed the pipeline and performed data preprocessing and analysis, with extensive and thorough guidance from AI, JC and VDC. RB drafted the manuscript, supplementary and all its figures. PAC offered detailed suggestions in introduction, interpretation of results in discussion on connectivity differences of psychosis biotypes. PAC revised the manuscript substantially, maintaining necessary results and discussion in the manuscript. PAC also processed necessary information for cross comparison of psychosis biotypes with B-SNIP study. KJ shared his work, on labeling 105 multiscale ICNs into domains and subdomains, which was relied upon to represent the results. KJ reviewed the manuscript and supported in interpretation of results, offering his inputs in the discussion. PB worked on analysis of low dimensional subspace and its associated components. PB performed analysis and generated results, figures for subspace characterization. CDC, JRB, JAT and ZF offered detailed feedback across the entire work and suggested inclusion of additional information to the work, which have been incorporated and revised accordingly. PAC, CDC, and JRB offered feedback and guidance on discussing the clinical implications of the results. Entire manuscript was thoroughly reviewed by all authors, suggesting necessary revisions. All authors have approved the submitted version of the manuscript and agree upon the integrity of the work presented.

## MATERIALS AND CORRESPONDENCE

Any correspondence regarding materials request to be directed to either Ram Ballem or Armin Iraji.

Name: Ram Ballem Postal Address: 55 Park Place NE 18^th^ Floor – TReNDS Center Atlanta, GA – 30303 Name: Armin Iraji Postal Address: 55 Park Place NE 18^th^ Floor – TReNDS Center Atlanta, GA – 30303 Phone: (+1) 404-413-4978

