## SUPPLEMENTARY MATERIALS for "Mapping the Psychosis Spectrum – Imaging Neurosubtypes from Multi-Scale Functional Network Connectivity"

**Section 1. Participants**

The data used in all analyses are from 2103 participants recruited by the Bipolar-Schizophrenia Network on Intermediate Phenotypes (B-SNIP 1 & 2) Consortium (1), collected from multiple sites. Storage, management, and access to the data used for this study are overseen by the National Institute of Mental Health (NIMH) through the National Data Archive. Participants under the B-SNIP1 and 2 were all given a full explanation of the study procedures and written consent was obtained. All sites followed a standardized procedure for recruitment and data acquisition across diagnostic, clinical, and cognitive assessment techniques (refer Supplementary Table 1 for imaging parameters and site information).

The cognitive assessment of participants was administered through the Brief Assessment of Cognition in Schizophrenia (BACS) (2). BACS offers a reliable cognitive evaluation of individuals, involving 6 different techniques that span across four cognitive domains: Verbal Memory, Working Memory, Processing Speed, Reasoning, and Problem Solving. Six different subtasks involve list learning, digit sequencing, token motor task, category instances task/controlled oral word association, symbol coding and executive function Tower of London. These subtasks are aggregated into one to represent the overall composite score, integrating performance across all four cognitive domains. All BACS scores for participants are normalized and stratified across age and sex (2) . Apart from BACS, the participants were also administered on Wechsler Memory Scale (3) for cognitive assessment of individuals with subtests capturing forward and backward memory scores. Participants with psychosis recruited under B-SNIP2 indicate categorical values regarding the questions such as difficulties in learning math, reading/writing, or repetition of grades, requiring summer school or special education.

Additionally, participants in the study underwent clinical interviews for DSM IV (4). Clinical characteristics of participants with psychosis and controls were both rated on Birchwood Social Functioning (BSFS) (5) and Global Assessment of Functioning (GAF) (6). Participants with psychosis were rated on the Montgomery-Asberg depression rating scale (MADRS) (7), Positive and Negative syndrome scale (8), Young Mania rating scale (YMRS) (9), Schizo-Bipolar Scale (SBS) (10) and Hollingshead Two-Factor Socioeconomic rating scale (SES) (11).

**Section 2. Image Acquisition and Preprocessing**

Participants of the study underwent a 5-minute-long resting state fMRI scan on 3T MRI scanner. During the scan, they were instructed to have their eyes open, focusing on a crosshair displayed on a monitor while remaining still. Additionally, a custom-built head coil cushion was employed to minimize head motion during the scan. Details regarding the acquisition parameters are mentioned in the Supplementary Table 1.

**Supplementary Table 1. Imaging Parameters of B-SNIP1 and B-SNIP2 multisite fMRI data**

| **Site** | **Acquisition Matrix (mm)** | **Voxel Size (mm)** | **Vendor** |
| --- | --- | --- | --- |
| **B-SNIP1** |  |  |  |
| Site-1 | 64 × 64 | 3.4 × 3.4 × 3 | Siemens Triotim |
| Site-2 | 65 × 64 | 3.4 × 3.4 × 5 | Siemens Allegra |
| Site-3 | 66 × 64 | 3.4 × 3.4 × 4 | Siemens Triotim |
| Site-4 | 67 × 64 | 3.4 × 3.4 × 4 | Philips |
| Site-5 | 68 × 64 | 3.4 × 3.4 × 4 | GE Signa HDX |
| **B-SNIP2** |  |  |  |
| Site-6 | 260 × 260 | 1.02 × 1.02 × 1.02 | GE HDx |
| Site-7 | 256 × 240 | 1 × 1 × 1.02 | Siemens Skyra |
| Site-8 | 260 × 260 | 1.02 × 1.02 × 1.02 | GE HDxt |
| Site-9 | 256 × 240 | 1 × 1 × 1.02 | Philips Achieva |
| Site-10 | 256 × 240 | 1 × 1 × 1.02 | Philips dSteam Achieva |

The preprocessing pipeline began by discarding the initial ten volumes to ensure steady state magnetization, eliminating unstable signals caused by radio frequency excitations. Rigid body motion correction was applied using FSL’s mcflirt function, aligning volumes to first volume. Distortion correction was performed with FSL’s applytopup function, followed by slice timing correction for temporal differences at each TR using SPM toolbox.

Scans were then warped to Montreal Neurological Institute (MNI) space using an echo planar imaging (EPI) template (12) and resampled to imaging data of 3 × 3 × 3 mm^3^ voxel space through SPM normalization tool. Finally, spatially smoothing was applied using a Gaussian kernel with a full-width half maximum of 6 mm to reduce noise and enhance the signal-to-noise ratio.

**Section 3. NeuroMark 2.2: A Replicable Data-Driven Multi-Scale Functional Atlas**

Subject-specific multi-scale intrinsic connectivity networks (ICNs) were estimated through independent component analysis spatially constrained to the NeuroMark 2.2 atlas of 105 highly reliable ICNs (13) (available at <https://trendscenter.org/data>; refer Supplementary Table 2).

**Supplementary Table 2. Domain, subdomain, and individual labels for the 105 multi-scale ICNs within the Neuromark 2.2 reference template**

| **Domain** | **ICN** | **Label** | **Peak Coordinates** | | |
| --- | --- | --- | --- | --- | --- |
|  |  |  | **x** | **y** | **z** |
| **Cerebellar Domain (CB)** | | | | | |
|  | 1 | anterior cerebellum (aCB) | 39 | -40 | -40 |
|  | 2 | posterior cerebellum (pCB) | 30 | -82 | -37 |
|  | 3 | anterior ventromedial cerebellum (avmCB) | 12 | -55 | -52 |
|  | 4 | ventromedial cerebellum (vmCB) | -21 | -55 | -52 |
|  | 5 | right cerebellum (rCB) | 30 | -55 | -43 |
|  | 6 | left cerebellum (lCB) | -27 | -58 | -40 |
|  | 7 | bilateral cerebellum (bCB) | 21 | -67 | -31 |
|  | 8 | medial cerebellum (mCB) | -3 | -61 | -31 |
|  | 9 | anterior dorsomedial cerebellum (admCB) | 0 | -52 | -19 |
|  | 10 | left anterior dorsomedial cerebellum (ladmCB) | -15 | -46 | -25 |
|  | 11 | right anterior dorsomedial cerebellum (radmCB) | 15 | -46 | -22 |
|  | 12 | vermis (Ver) | 0 | -49 | -13 |
|  | 13 | dorsomedial cerebellum (dmCB) | -3 | -64 | -16 |
| **Visual Domain (VI)** | | | | | |
| *Occipitotemporal Subdomain (OT)* | 14 | fusiform gyrus (FG) | 33 | -46 | -16 |
|  | 15 | ventromedial occipitotemporal cortex (vmOTC) | 30 | -49 | -10 |
|  | 16 | left medial occipitotemporal cortex (lmOTC) | -21 | -49 | -7 |
|  | 17 | right medial occipitotemporal cortex (rmOTC) | 21 | -46 | -7 |
|  | 18 | anterior medial occipitotemporal cortex (amOTC) | 15 | -67 | 11 |
|  | 19 | occipitotemporal junction (OTJ) | 45 | -61 | 5 |
| *Occipital Subdomain (OC)* | 20 | extrastriate cortex (ESC) | -42 | -73 | -4 |
|  | 21 | left extrastriate cortex (lESC) | -27 | -76 | -4 |
|  | 22 | right extrastriate cortex (rESC) | 30 | -73 | -1 |
|  | 23 | striate cortex (Str) | -6 | -88 | -4 |
|  | 24 | cuneus (CUN) | -9 | -94 | 26 |
|  | 25 | occipital pole (OP) | 27 | -94 | -4 |
| **Paralimbic Domain (PL)** | | | | | |
|  | 26 | lateral temporal pole (latTP) | 51 | 14 | -25 |
|  | 27 | left temporal pole (lTP) | -30 | 2 | -37 |
|  | 28 | right temporal pole (rTP) | 27 | 5 | -37 |
|  | 29 | left medial temporal cortex (lmTC) | -21 | -7 | -25 |
|  | 30 | left medial temporal pole (lmTP) | -24 | 8 | -25 |
|  | 31 | right medial temporal cortex (rmTC) | 21 | -7 | -25 |
|  | 32 | right medial temporal pole (rmTP) | 27 | 8 | -25 |
|  | 33 | bilateral medial temporal pole (bmTP) | -24 | 5 | -28 |
|  | 34 | bilateral temporal pole (bTP) | 27 | 8 | -34 |
|  | 35 | entorhinal cortex (EC) | 24 | 2 | -34 |
|  | 36 | hippocampal–entorhinal complex (HEC) | 24 | -13 | -19 |
| **Subcortical Domain (SC)** | | | | | |
| *Extended Hippocampal Subdomain (EH)* | 37 | right hippocampus/parahippocampal cortex (rHPC) | 30 | -31 | -4 |
|  | 38 | left hippocampus/parahippocampal cortex (lHPC) | -30 | -34 | -4 |
|  | 39 | thalamus/hippocampus/amygdala (Thal-Amg) | 9 | -31 | 5 |
| *Extended Thalamic Subdomain (ET)* | 40 | thalamus/hippocampus (Thal-Hip) | -3 | -13 | 2 |
|  | 41 | thalamus (Thal) | 9 | -7 | 8 |
|  | 42 | diencephalon/midbrain (DiMid) | 6 | -13 | -4 |
|  | 43 | anterior diencephalon (aDi) | 6 | -4 | -4 |
|  | 44 | right thalamus (rThal) | 21 | -13 | -1 |
|  | 45 | left thalamus (lThal) | -21 | -13 | -1 |
| *Basal Ganglia Subdomain (BG)* | 46 | dorsal striatum/thalamus (DS-Thal) | -21 | 5 | 5 |
|  | 47 | right lentiform nucleus (rLN) | 27 | 5 | -4 |
|  | 48 | left lentiform nucleus (lLN) | -27 | 2 | -4 |
|  | 49 | lentiform nucleus (LN) | 24 | 8 | -1 |
|  | 50 | left caudate (lCaud) | -21 | 17 | 5 |
|  | 51 | right caudate (rCaud) | 21 | 17 | 5 |
|  | 52 | left basal ganglia (lBG) | -18 | 11 | -10 |
|  | 53 | right basal ganglia (rBG) | 18 | 14 | -7 |
|  | 54 | bilateral basal ganglia (bBG) | 9 | 8 | -7 |
| **Sensorimotor Domain (SM)** | | | | | |
|  | 55 | supplementary motor area (SMA) | 0 | -1 | 59 |
|  | 56 | medial supplementary motor area (mSMA) | -9 | 2 | 44 |
|  | 57 | posterior supplementary motor area (pSMA) | -15 | -10 | 62 |
|  | 58 | precentral gyrus (PrCG) | 36 | -10 | 41 |
|  | 59 | superior sensorimotor cortex (sSMC) | 21 | -25 | 59 |
|  | 60 | paracentral lobule (PCL) | 0 | -25 | 65 |
|  | 61 | right superior postcentral gyrus (rsPoCG) | 39 | -22 | 59 |
|  | 62 | left superior postcentral gyrus (lsPoCG) | -39 | -22 | 62 |
|  | 63 | superior parietal lobule (SPL) | 21 | -52 | 71 |
|  | 64 | inferior postcentral gyrus (iPoCG) | -51 | -10 | 32 |
|  | 65 | supramarginal gyrus (SMG) | 54 | -22 | 29 |
|  | 66 | posterior postcentral gyrus (pPoCG) | -54 | -25 | 38 |
|  | 67 | anterior inferior parietal lobe (aIPL) | 60 | -25 | 41 |
|  | 68 | posterior inferior parietal lobe (pIPL) | -60 | -40 | 41 |
| **Higher Cognition Domain (HC)** | | | | | |
| *Insular Temporal Subdomain (IT)* | 69 | left posterior insular cortex (lPIC) | -39 | -7 | 17 |
|  | 70 | right posterior insular cortex (rPIC) | 36 | -13 | 11 |
|  | 71 | dorsal anterior insular cortex (dAIC) | -36 | 5 | 11 |
|  | 72 | ventral anterior insular cortex (vAIC) | -42 | -1 | -10 |
|  | 73 | medial ventral anterior insular cortex (mvAIC) | 42 | 5 | -13 |
|  | 74 | posterior superior temporal gyrus (pSTG) | -39 | -31 | 14 |
|  | 75 | superior temporal gyrus (STG) | 63 | -25 | 5 |
| *Temporoparietal Subdomain (TP)* | 76 | left middle temporal gyrus/temporoparietal junction (lMTG-TPJ) | -48 | -46 | 8 |
|  | 77 | right middle temporal gyrus/temporoparietal junction (rMTG-TPJ) | 48 | -43 | 8 |
|  | 78 | bilateral temporoparietal junction/social mind areas (bTPJ-S) | 57 | -46 | 23 |
|  | 79 | posterior temporal cortex (pTC) | 51 | -46 | 14 |
|  | 80 | inferior posterior temporal cortex (ipTC) | -51 | -46 | -7 |
| *Frontal Subdomain (FR)* | 81 | right inferior frontal gyrus (rIFG) | 51 | 20 | 17 |
|  | 82 | left inferior frontal gyrus (lIFG) | -48 | 26 | 5 |
|  | 83 | ventrolateral prefrontal cortex (VLPFC) | 51 | 23 | 17 |
|  | 84 | inferior left ventrolateral prefrontal cortex (ilVLPFC) | -48 | 20 | 23 |
|  | 85 | left ventrolateral prefrontal cortex (lVLPFC) | -45 | 20 | 23 |
|  | 86 | posterior inferior frontal gyrus (pIFG) | -36 | 14 | 29 |
|  | 87 | orbitofrontal cortex (OFC) | -24 | 50 | -7 |
|  | 88 | frontal pole (FP) | -30 | 62 | 8 |
|  | 89 | Frontal eye field area (FEF) | -15 | 11 | 53 |
|  | 90 | anterior supplemental motor area (aSMA) | -9 | 8 | 62 |
| **Triple Network Domain (TN)** | | | | | |
| *Central Executive Network Subdomain (CE)* | 91 | right inferior parietal lobe/dorsolateral prefrontal cortex (rIPL-rDLPFC) | 48 | -58 | 47 |
|  | 92 | left inferior parietal lobe/dorsolateral prefrontal cortex (lIPL-lDLPFC) | -45 | -61 | 50 |
|  | 93 | bilateral inferior parietal lobe/dorsolateral prefrontal cortex (bIPL-lDLPFC) | -51 | -58 | 41 |
| *Default Network Subdomain (DN)* | 94 | right inferior parietal lobe/posterior cingulate cortex (rIPL-rPCC) | 48 | -64 | 38 |
|  | 95 | left inferior parietal lobe/posterior cingulate cortex (lIPL-lPCC) | -48 | -64 | 35 |
|  | 96 | posterior cingulate cortex (PCC) | -12 | -58 | 20 |
|  | 97 | ventral posterior cingulate cortex (vPCC) | -12 | -52 | 11 |
|  | 98 | ventral precuneus (vPRCU) | -9 | -67 | 35 |
|  | 99 | dorsal precuneus (dPRCU) | -6 | -55 | 50 |
|  | 100 | dorsomedial prefrontal cortex (dmPFC) | 0 | 47 | 35 |
|  | 101 | medial prefrontal cortex (mPFC) | 0 | 50 | 20 |
| *Salience Network Subdomain (SN)* | 102 | anterior cingulate cortex (ACC) | 0 | 41 | 2 |
|  | 103 | dorsal anterior cingulate cortex (dACC) | 0 | 35 | 11 |
|  | 104 | anterior cingulate cortex/anterior insula (ACCAI) | 0 | 32 | 20 |
|  | 105 | anterior insula/anterior cingulate cortex (AIACC) | 33 | 23 | -4 |

**Section 4. Subject-Specific Multi-Scale Functional Network Connectivity (msFNC)**

Time courses were detrended by removing linear, quadratic, and cubic trends. Motion was further addressed by performing a regression of time courses with the six motion realignment parameters and their derivatives. Outliers were detected based on the median absolute deviation, spike threshold (c1) = 2.5 (14) and replaced with the best estimate using a third-order spline fit to the clean portions of the time courses. Lastly, bandpass filtering was applied using a fifth-order Butterworth filter with a cutoff frequency of 0.01 Hz - 0.15 Hz.

**Section 5. Analysis Datasets: Discovery and Replication Data**

Across the entire dataset, the age of controls and psychosis participants statistically did not show any difference (p-value = 0.14)**.** However, the chi-squared test statistic rejects the null hypothesis that sex and diagnoses (controls or psychosis) are independent (p-value < 0.001). Comparing controls with DSM groups, age was statistically significant between controls - SAD (p-value < 0.01), however controls - SZ (p-value = 0.49) and controls - BP was not significant (p-value = 0.52). The SZ and SAD DSM groups were also statistically significant between each other (p-value < 0.05) as well as BP - SAD (p-value < 0.01), but SZ - BP didn’t show a significant difference (p-value = 0.23).

Within the discovery and replication sets, age distributions of controls and psychosis were not statistically significant (discovery set control-psychosis age statistics: t=-0.96, p=0.33, df=1389, effect=-0.65; replication set control-psychosis age statistics: t=-1.32, p=0.18, df=360, effect=-1.78). A Pearson chi-squared test indicated that within the discovery set, sex and psychosis were not independent (chi-squared test statistic = 12.22, p < 0.001), while in the replication set, they were independent (chi-squared test statistic = 0.25, p = 0.38). Additionally, the division also ensured that psychosis participants with chlorpromazine equivalent information existed in both these datasets (N=351/891, 40% of the psychosis affected in the discovery set).

**Section 6. Low-Dimensional Latent Network Connectivity Subspace**

Extraction of the Latent Network Connectivity subspace (LNC) from msFNC began by applying ICA on the msFNC of the discovery set. The number of independent components to extract was determined through a prior PCA step. After performing PCA on the msFNC of the discovery set, we examined the cumulative variance explained by the principal components. This variance curve showed an elbow at the 3^rd^ principal component (Supplementary Fig.1), till which 60% of msFNC variance was contained, guiding our decision to extract three independent components.

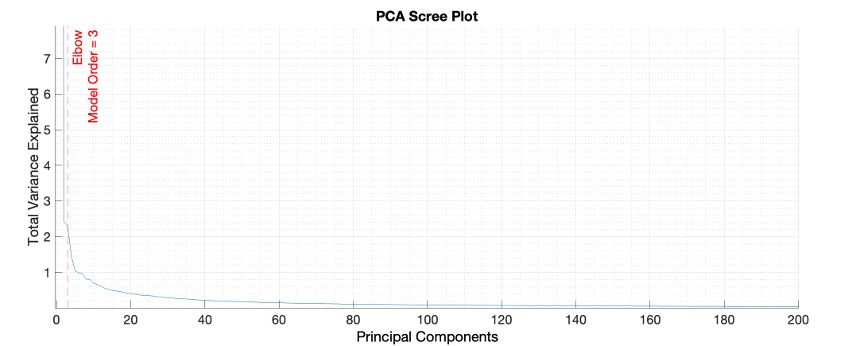

**Supplementary Fig.1**: **Multi-scale Functional Network Connectivity Principal Component Analysis (PCA) Scree Plot.** Variance explained by the Principal Components obtained from applying PCA on the msFNC of the discovery set. The observed elbow of the variance is explained in the scree plot at Principal Component Number 3.

**Section 6.1. Cognitive Related Component**

The first independent component, LNC component 1 was extracted from the msFNC of all participants in the discovery set, including both controls and psychosis (Fig.3D in the main manuscript). The component highlights connectivity in HC, CB, SM, and TN domains. Specific subdomains (IT and FR) overlap spatially with regions (e.g., superior temporal gyrus/STG, inferior frontal gyrus, and dorsolateral prefrontal cortex) crucial for language processing, working memory, and attention (15). These regions correspond to cognitive domains assessed by BACS (16). The Latent Network Connectivity Projections (LNC Projections 1) of this LNC component 1 showed a significant association with cognitive assessment scores (Supplementary Fig.2). Notably, BACS Composite (Supplementary Fig.2A), Digit Sequencing (Supplementary Fig.2B), Tower of London (Supplementary Fig.2C), and both WMS subtests - Forward and Backward (Supplementary Fig.2D&E) displayed associations in both discovery and replication sets. This led us to identify LNC component 1 as the cognitive related component.

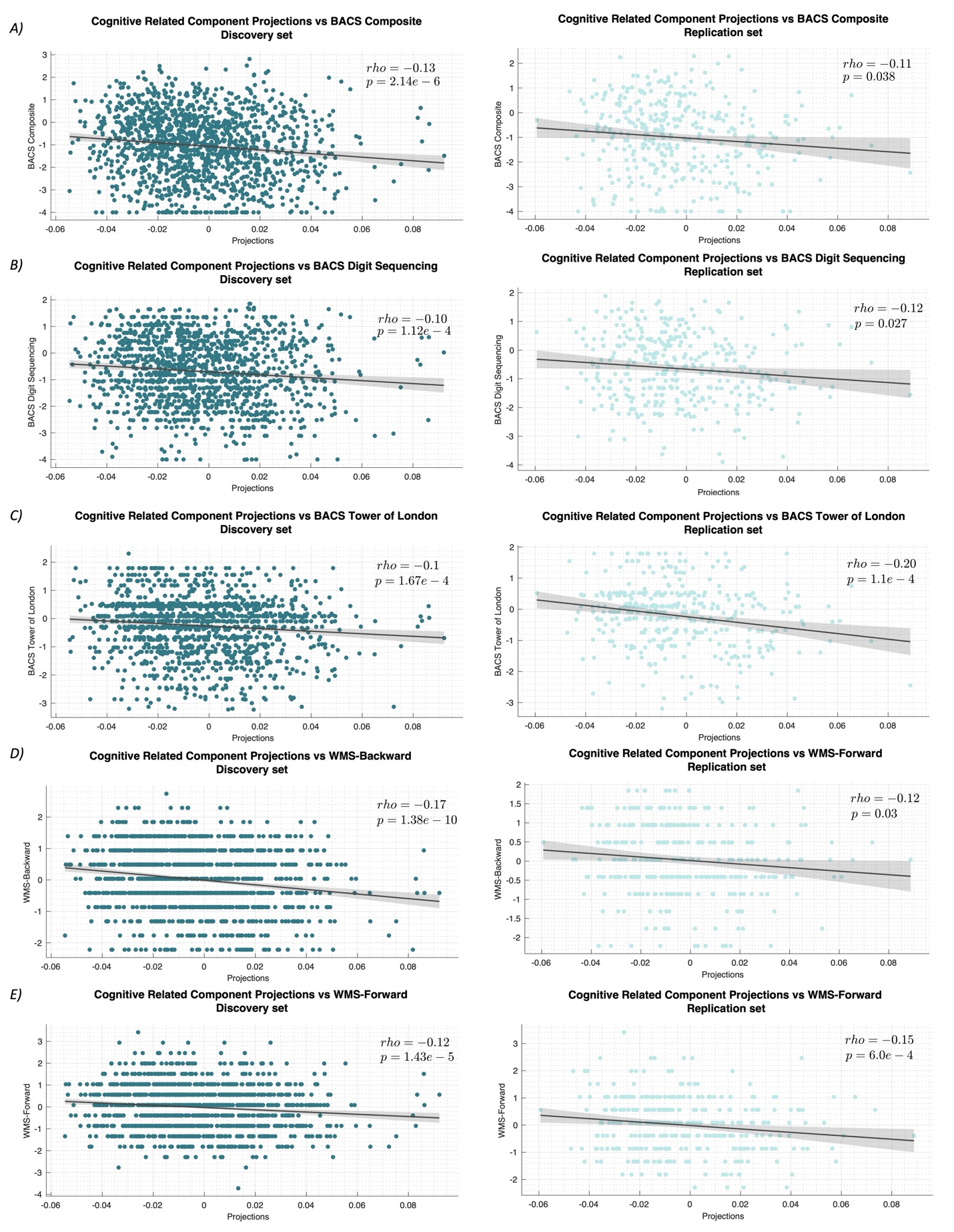

**Supplementary Fig.2**: **Cognitive assessment scores associations with Cognitive related component projections in discovery and replication sets.** A) BACS composite score association with cognitive related component projections in discovery and replication sets. B) BACS Digit Sequencing assessment score associations with cognitive related component projections in discovery and replication sets. C) BACS Tower of London assessment score associations with cognitive related component projections in discovery and replication sets. D) WMS Forward subtest assessment score associations with cognitive related component projections in discovery and replication sets. E) WMS Backward subtest assessment score associations with cognitive related component projections in discovery and replication sets. All associations above are statistically significant and replicable.

**Section 6.2. Typical Component**

Like the first component, the second independent component, LNC 2, was extracted from the msFNC of all participants in the discovery set, including both controls and psychosis (Fig.3E in the main manuscript). This independent component resembles a modular pattern that favored within domain ICN connectivity over between domain connections, a typical pattern observed in average functional network connectivity of groups like control or psychosis (Supplementary Fig.3A&B). Since LNC component 2 reflects the central tendency or population level property, we named LNC 2 as the typical component. It reflects a modular brain organization with strong within-domain connectivity and weak between-domain connectivity, consistent with established brain organization features (17).

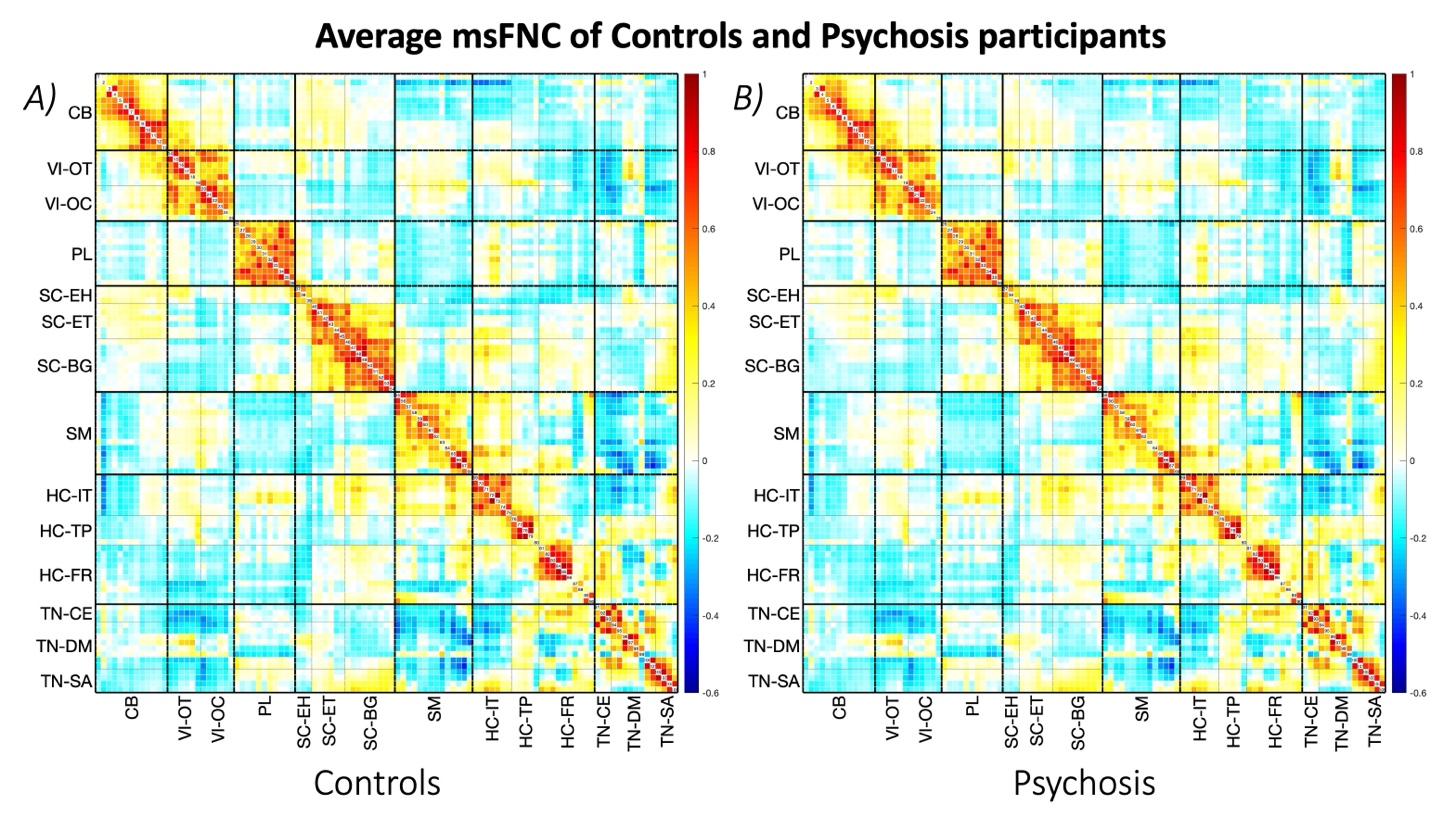

**Supplementary Fig.3**: **Average Multi-scale Functional Network Connectivity (msFNC) of Controls (CON) and Psychosis participants (PSY) in the discovery set.** msFNC represents connectivity between 105 Multi-scale Intrinsic Connectivity Networks (msICNs). Refer Fig.1 and Supplementary Table 2 for additional information on the 105 multi-scale intrinsic connectivity networks. A) Average msFNC of the CON group in the discovery set. B) Average msFNC of the PSY group in the discovery set.

**Section 6.3. Psychosis Related Component**

Like LNC 1 and 2, the third independent component, LNC 3, was extracted from the msFNC of all participants in the discovery set, including both controls and psychosis (Fig.3F in the main manuscript). This independent component resembles key connectivity patterns in CB, SM, VI, SC, and HC regions, characteristic of psychosis, aligning with group differences between controls and psychosis participants (18–20) (Supplementary Fig.4A&B). Since LNC component 3 captures this pattern of aberrant connectivity associated with psychosis, we named LNC 3 as a psychosis related component.

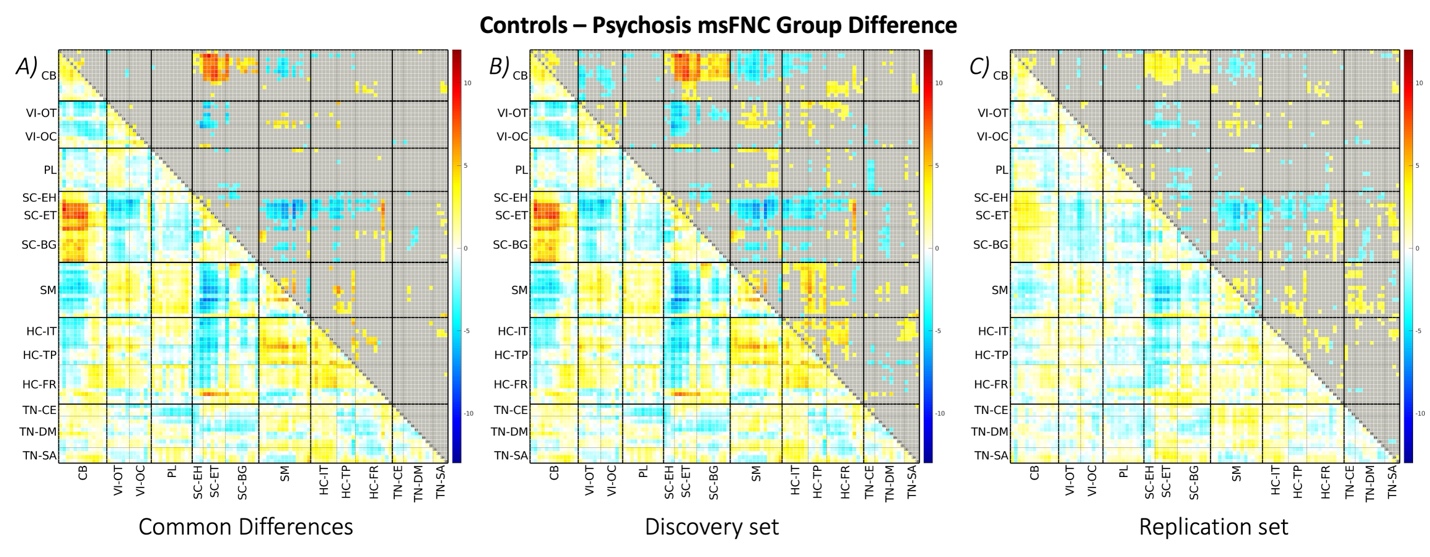

**Supplementary Fig.4**: **Group Differences between Controls (CON) - Psychosis (PSY) groups along Multi-scale Functional Network Connectivity (msFNC).** msFNC represents connectivity between 105 Multi-scale Intrinsic Connectivity Networks (msICNs). Refer Fig.1 and Supplementary Table 2 for additional information on the 105 multi-scale intrinsic connectivity networks. These 105 × 105 matrices display the t-value of discovery set in the lower triangle, where each cell reflects group differences in connectivity between two msICNs. The upper triangle shows the statistical significance using -log(q-value).*sign(t-value) for the discovery set and -log(p-value).*sign(t-value) for replication set, with non-significant ones grayed out. The t-value ranges from -13 to 12. A) msFNC Group difference matrix between the CON and PSY groups, common significant results of discovery set and replication set. B) msFNC Group difference matrix between CON and PSY group significant in the discovery set. C) msFNC Group difference matrix between CON and PSY group significant in the replication set.

**Section 7. Psychosis Imaging Neurosubtypes**

To identify Psychosis Imaging Neurosubtypes (PINs), a preliminary step was to estimate the optimal number of clusters within all the LNC projections of psychosis participants of discovery set. This was done using various cluster evaluation methods available within MATLAB, including the Calinski-Harabasz index, silhouette evaluation, and cluster dispersion ratio. All these methods consistently showed that the optimal number of clusters was 3 (Supplementary Fig.5A-C).

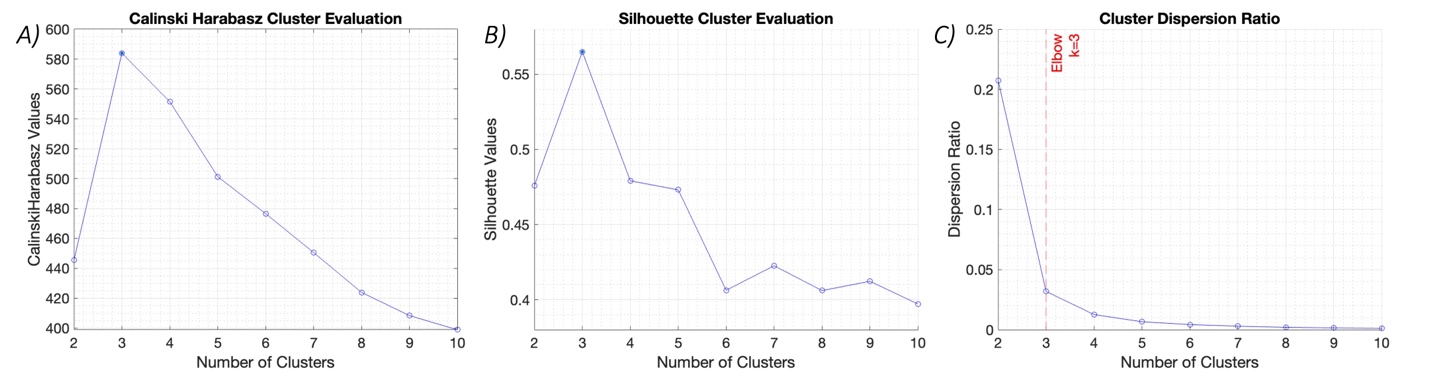

**Supplementary Fig.5**: **Psychosis Imaging Neurosubtypes cluster evaluations** A-C) Cluster evaluation of the psychosis group Latent Network Connectivity projections in the discovery dataset over 2 - 10 clusters. A) Calinski Harabasz Index plot explaining cluster number vs. CH index, revealing the highest index value at k=3. B) Silhouette plot of cluster numbers 2-10 vs. silhouette value at each cluster number, revealing the highest silhouette value at k=3. C) Cluster dispersion ratio plot, with cluster numbers 2-10 vs. their dispersion ratio of within cluster distances to between cluster distances. The elbow of this plot was observed at k=3.

The K-means clustering algorithm was employed to identify clusters, with parameters including the squared Euclidean distance metric, a maximum of 10^8^ iterations, and 50 replications to ensure the algorithm’s convergence towards an optimal solution. Centroid initialization was performed using the k-means++ method, favoring efficiency of convergence. Participants in the hold-out replication set were subsequently assigned to these clusters by comparing their LNC projections to PINs centroids, based on minimum squared Euclidean distance, calculated via MATLAB’s pdist2 function.

PINs showed statistically significant age differences with controls in discovery dataset (control vs. PIN-1 t=-3.99, p<0.001, df=755, effect=-3.75; control vs. PIN-2 t=-2.81, p<0.01, df=685, effect=-2.91; control vs. PIN-3 t=2.65, p<0.01, df=945, effect=2.06). These significant age differences were observed to be replicated between control and PIN-2 (control vs. PIN-1 t=-1.85, p=0.06, df=182, effect=-3.69; control vs. PIN-2 t=-2.94, p<0.01, df=174, effect=-6.27; control vs. PIN-3 t=0.54, p=0.59, df=252, effect=0.82).

PINs also exhibited a statistically significant age difference between themselves in the discovery set (PIN-1 vs. PIN-2 t=0.72, p=0.47, df=442, effect=0.84; PIN-1 vs. PIN-3 t=6.23, p<0.001, df=702, effect=5.83; PIN-2 vs. PIN-3 t=4.89, p<0.001, df=632, effect=4.99). Replicability of this age difference was observed between PIN-1 and PIN-3, PIN-2 and PIN-3 in the replication set (PIN-1 vs. PIN-2 t=-1.13, p=0.26, df=106, effect=-2.58; PIN-1 vs. PIN-3 t=2.52, p<0.05, df=184, effect=4.53; PIN-2 vs. PIN-3 t=3.70, p=<0.001, df=176, effect=7.1).

The Chi-squared test statistic indicated that sex and the identified PINs (controls, PIN-1, PIN-2, PIN-3) were independent (chi-squared test statistic = 13.09, p<0.01). Similarly, the replication set identified that the sex and neurosubtypes were independent (chi-squared test statistic=3.08, p=0.61).

The significant differences in age and sex between controls and PINs, and between PINs, may have influenced the overall analysis. Consequently, covariates such as age, sex, race, site, and head motion (where applicable) were included as regressors in all analyses characterizing neurosubtypes. Despite adjusting for these covariates, the identification of neurosubtypes and their subsequent characterizations were also conducted using the cleaned msFNC data (i.e., the residual msFNC after correcting for the covariates in a GLM). The clustering results showed that neurosubtype labels were approximately 90% consistent across analyses.

**Section 8. Psychosis Imaging Neurosubtypes - Characterization and Validation**

**Section 8.1. Psychosis Imaging Neurosubtypes Cognitive Characterization**

Statistically significant differences in cognition were observed between the PINs and controls in the replication set (represented with an *, p < 0.05, in Supplementary Fig.6A). Exceptions include the Tower of London assessment score and WMS Forward subtest (WMS-F), which did not show statistical significance between controls and PIN-2, controls and PIN-3.

Differences in cognitive assessments among the PINs themselves are presented in Supplementary Fig.6B. In the replication set, PIN-1 displayed significant differences from PIN-2 along the BACS composite score, verbal memory, digit sequencing, and tower of London. PIN-1 also showed a significant difference from PIN-3 in the composite score and tower of London.

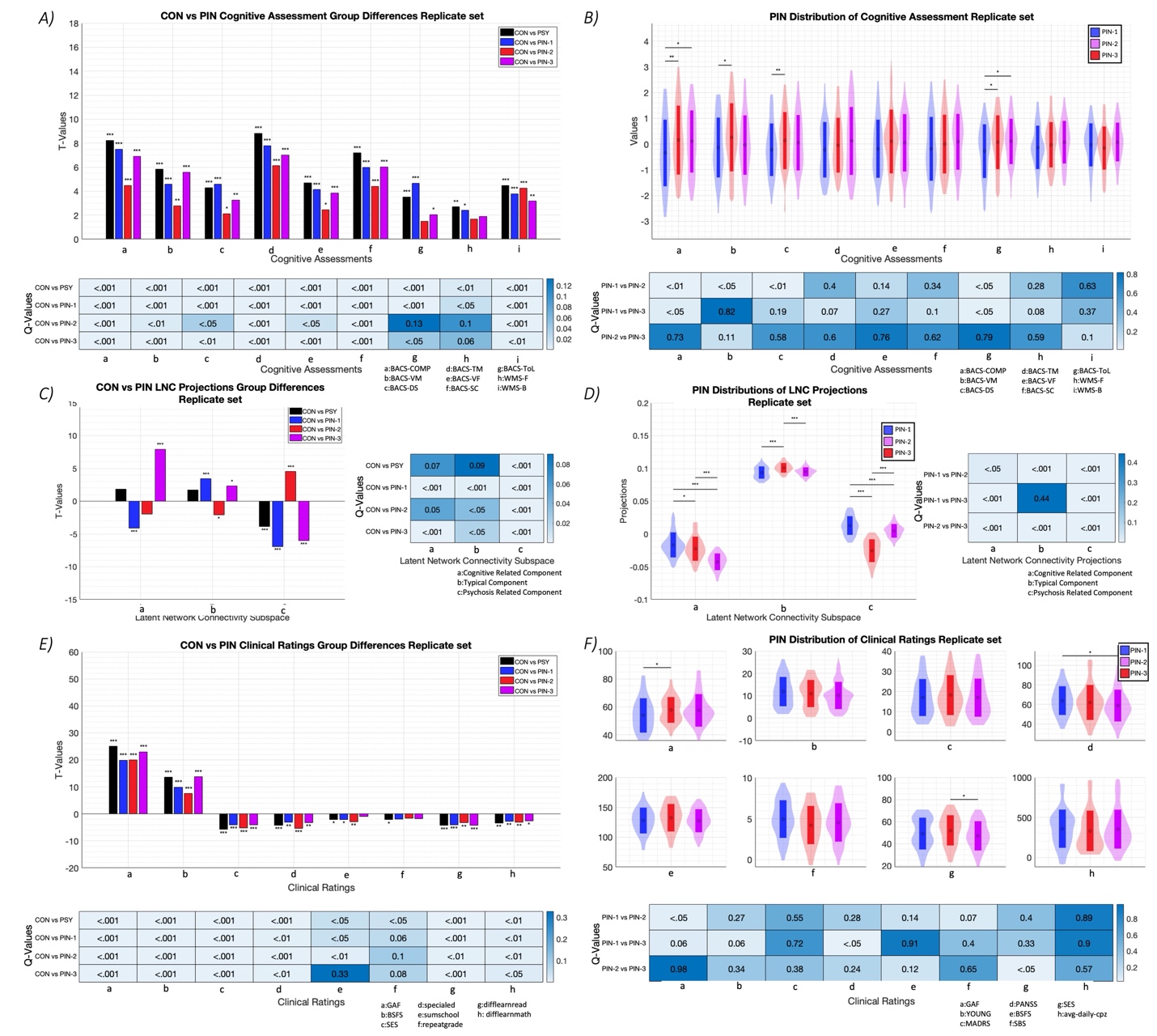

**Supplementary Fig.6**: **Group Differences between the Control (CON) - Psychosis Imaging Neurosubtypes (PINs) and within PINs across Cognitive Assessment Scores (Brief Assessment of Cognition in Schizophrenia, BACS and Weschler Memory Scale, WMS), Latent Network Connectivity (LNC) Projections, and Clinical Variables, in Replication set.** A,B) Different cognitive assessment scores involve BACS Composite Score(BACS-COMP), Verbal Memory (BACS-VM), Token Motor (BACS-TM), Verbal Fluency (BACS-VF), Symbol Coding (BACS-SC), Tower of London (BACS-ToL) and WMS Forward (WMS-F) and WMS Backward (WMS-B) subtests. A) Bar plots indicate cognitive assessment scores group differences in the replication set, the x-axis representing different cognitive assessments, and the y-axis showing the T-Values (t-value ranges from 0 to 18) of the group differences. The heatmap below indicates the P-Values of group differences in cognitive assessment scores, between the CON and PINs of the replication set. B) Violin plots represent PINs distributions of cognitive assessment scores in the replication set, with error bars indicating the mean and standard deviation. The heatmap below indicates the P-Values for group differences in cognitive assessment scores, within neurosubtypes in the replication set. C,D) Different LNC projections involve Cognitive Associated Component projections (LNC projections 1), Typical Component projections (LNC projections 2), and Psychosis Related Component projections (LNC projections 3). C) Bar plots indicate different LNC projection group differences in the replication set, with the x-axis representing different LNC component projections and the y-axis showing the T-Values (ranging from -15 to 15) of the group difference. The heatmap adjacent to it indicates the P-Values of the group differences in LNC projections, between CON and PINs of the replication set. D) Violin plots represent the PINs distributions of LNC projections in the replication set, with error bars indicating the mean and standard deviation. The heatmap adjacent to it indicates the P-Values of the group difference in LNC projections within neurosubtypes of the replication set. E,F) Different clinical variables involve Global Assessment of Functioning (GAF), Young Mania Rating Scale (YMRS), Montgomery-Asberg Depression Rating Scale (MADRS), Positive and Negative Syndrome Scale (PANSS), Birchwood Social Functioning (BSFS), Schizo-Bipolar Scale (SBS), Socioeconomic Rating Scale (SES), and Average Daily Chlorpromazine Equivalents (avg-daily-cpz). E) Bar plots indicate group differences across clinical variables in the replication set, with the x-axis representing different clinical variables and the y-axis showing the T-Values (ranging from -20 to 60) of the group differences. The heatmap below indicates the P-Values of the group differences in clinical variables between CON and PINs in the replication set. F) Violin plots represent PINs distributions of clinical variables in the replication set, with error bars indicating the mean and standard deviation. The heatmap below indicates the P-Values of group differences in clinical variables within neurosubtypes of the replication set. A,C,E) Plots indicating group differences between CON and PINs using color bars to show different group comparisons (black = CON vs. PSY group, blue = CON vs. PIN-1, red = CON vs. PIN-2, and violet = CON vs. PIN-3). B,D,F) Plots indicating distributions within PINs (blue, red, and violet indicate PIN-1, PIN-2, and PIN-3 distributions respectively). A-F) In all the plots, a single * represents statistical significance (p-value < 0.05), with double ** representing a p-value < 0.01, and triple *** representing a p-value < 0.001. List of abbreviations – CON: Controls; PSY: Psychosis; LNC: Latent Network Connectivity; PIN: Psychosis Imaging Neurosubtype; PIN-1: Psychosis Imaging Neurosubtype-1; PIN-2: Psychosis Imaging Neurosubtype-2; PIN-3: Psychosis Imaging Neurosubtype-3; BACS: Brief Assessment of Cognition; BACS-COMP: Composite Score; BACS-VM: Verbal Memory; BACS-DS: Digit Sequencing; BACS-TM: Token Motor; BACS-VF: Verbal Fluency; BACS-ToL: Tower of London; WMS-F: Wechsler Memory Scale; WMS-F: Forward Subtest; WMS-B: Backward Subtest; GAF – Global Assessment of Function; YMRS: Young Mania Rating Scale; MADRS: Montgomery-Asberg Depression Rating Scale; PANSS: Positive and Negative Syndrome Scale; BSFS: Birchwood Social Functioning; SBS: Schizo-Bipolar Scale; SES: Socioeconomic Rating Scale; avg-daily-cpz: Average Daily Chlorpromazine Equivalents

**Section 8.2. Psychosis Imaging Neurosubtypes LNC Projections Characterization**

Controls exhibited statistically significant group differences in comparison to all PINs in the replication set (Supplementary Fig.6C), except between controls and PIN-2 along the cognitive related component projections (LNC 1 projections).

In the replication set, all PINs displayed statistically significant group differences between each other, except for PIN-1 and PIN-3 not showing significant differences along the typical component projections (Supplementary Fig.6D).

**Section 8.3. Psychosis Imaging Neurosubtypes Clinical Characterization**

PINs presented statistically significant differences comparing with the control group across all clinical variables in the replication set. However, the differences in educational difficulties such as summer school and the repetition of grades were not significant.

Psychosis PIN-1 and PIN-2 showed statistically significant differences in replication set along GAF score. PIN-1 and PIN-3 showed statistically significant differences in PANSS total score. PIN-2 and PIN-3 showed a significant difference in SES score (Supplementary Fig.6F)

Clinical group differences were also tested, by not accounting for multiple comparisons corrections. When FDR correction was not applied (Supplementary Fig.7), some significant differences were noted in discovery set, although most of them were not replicable. Specifically, PIN-2 showed a significant difference from PIN-3 along the YMRS scale in the discovery set. PIN-2 also differed significantly from PIN-1 on SES in the discovery set.

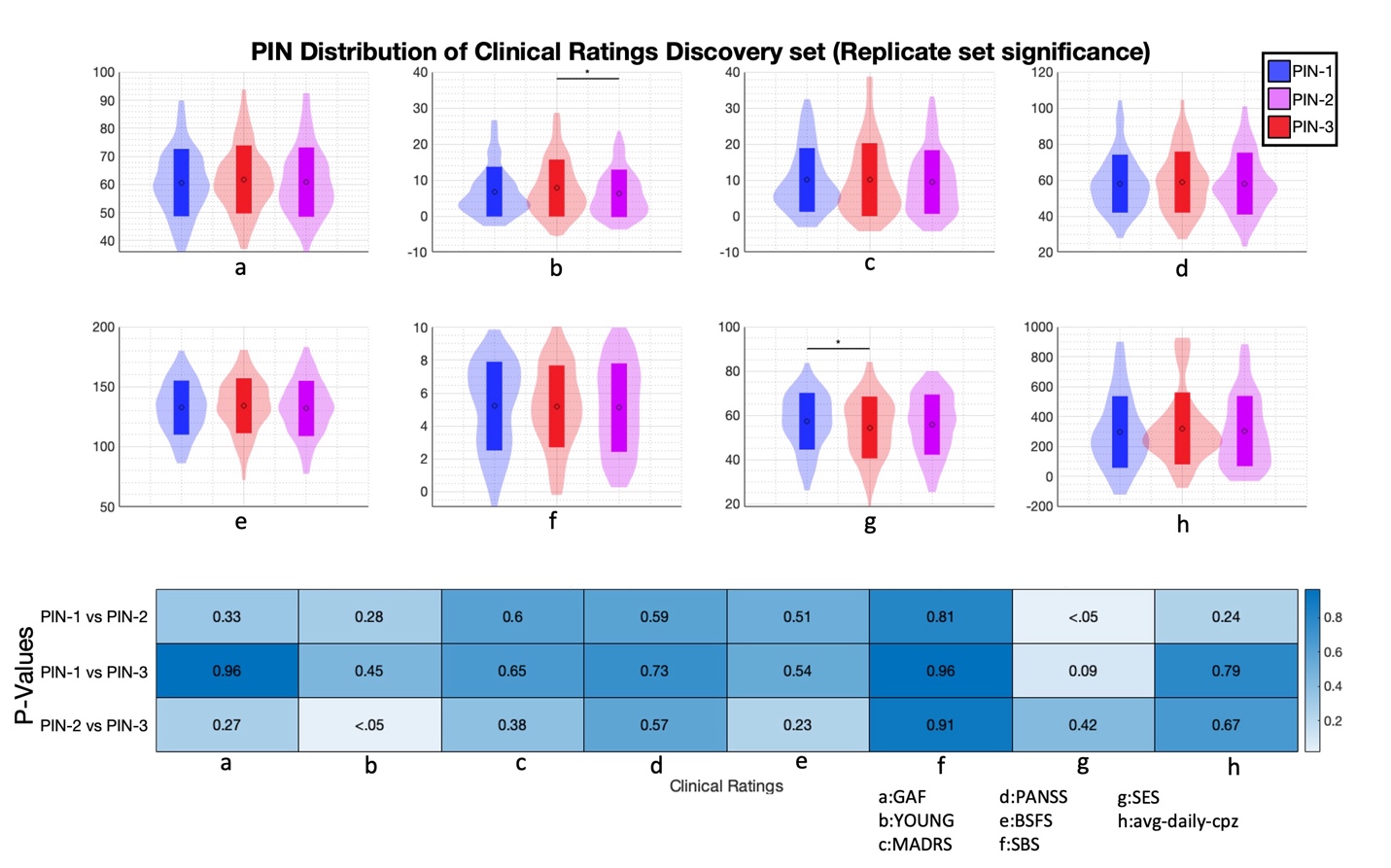

**Supplementary Fig.7**: **Group Differences within Psychosis Imaging Neurosubtypes (PINs) across Clinical Variables in the Discovery set (No FDR Correction) and Replication sets.** Different clinical variables involve Global Assessment of Functioning (GAF), Young Mania Rating Scale (YMRS), Montgomery-Asberg Depression Rating Scale (MADRS), Positive and Negative Syndrome Scale (PANSS), Birchwood Social Functioning (BSFS), Schizo-Bipolar Scale (SBS), Socioeconomic Rating Scale (SES), and Average Daily Chlorpromazine Equivalents (avg-daily-cpz). Violin plots representing the PINs distributions of clinical variables in the discovery set, with error bars indicating the mean and standard deviation. The plot uses blue, red and violet to indicate PIN-1, PIN-2 and PIN-3 respectively. Heatmap indicating P-Values of group difference in clinical variables within neurosubtypes of the discovery set, with no FDR correction. In all the plots, a single * represents statistical significance (p-value < 0.05), with double ** representing a p-value < 0.01, and triple *** representing a p-value < 0.001. List of abbreviations – CON: Controls; PSY: Psychosis; PIN: Psychosis Imaging Neurosubtype; PIN-1: Psychosis Imaging Neurosubtype-1; PIN-2: Psychosis Imaging Neurosubtype-2; PIN-3: Psychosis Imaging Neurosubtype-3; GAF: Global Assessment of Function; YMRS: Young Mania Rating Scale; MADRS: Montgomery-Asberg Depression Rating Scale; PANSS: Positive and Negative Syndrome Scale; BSFS: Birchwood Social Functioning; SBS: Schizo-Bipolar Scale; SES: Socioeconomic Rating Scale; avg-daily-cpz: Average Daily Chlorpromazine Equivalents

**Section 8.4. Psychosis Imaging Neurosubtypes msFNC Characterization**

Group differences in msFNC between controls and PINs, as well as differences within PINs, are shown in Supplementary Fig.8&9. Statistically significant differences in the discovery set between controls and neurosubtypes are highlighted in Supplementary Fig.8B,E&H, while differences between neurosubtypes are shown in Supplementary Fig.9B,E&H. Replicability results in the replication set are shown between controls and neurosubtypes in Supplementary Fig.8C,F&I and between neurosubtypes in Supplementary Fig.9C,F&I. Common differences found in both discovery and replication sets, indicating replicable group differences are shown in Supplementary Fig.8A,D&G for controls and neurosubtypes, and Supplementary Fig.9A,D&G for neurosubtype comparisons. The summary below comments on significant msFNC differences, detailing variations both within and between domains.

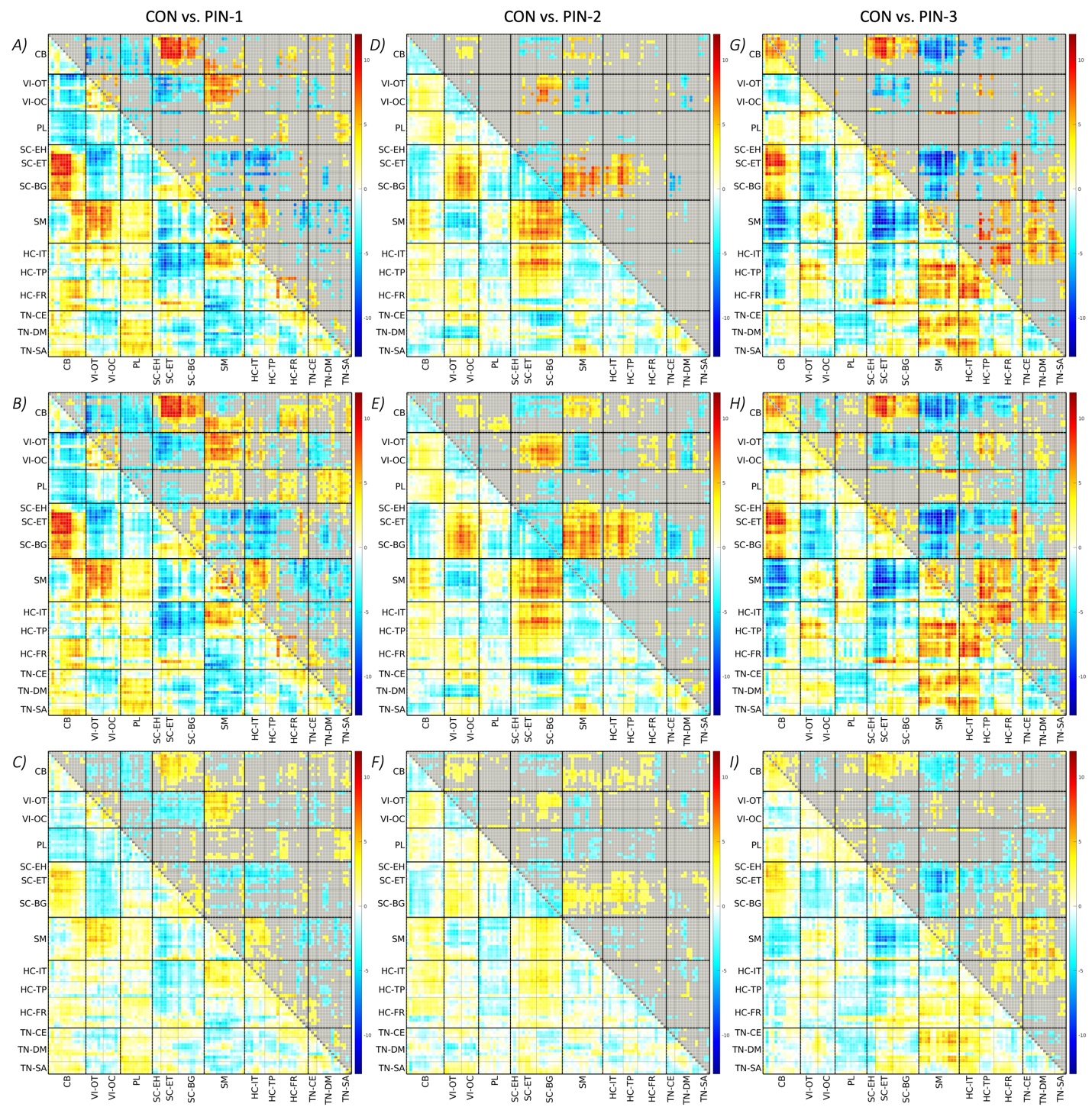

**Supplementary Fig.8**: **Group Differences between Controls (CON) - Psychosis Imaging Neurosubtypes (PINs) along Multi-scale Functional Network Connectivity (msFNC) in Discovery and Replication sets.** msFNC represents connectivity between 105 Multi-scale Intrinsic Connectivity Networks (msICNs). Refer Fig.1 and Supplementary Table 2 for additional information on the 105 multi-scale intrinsic connectivity networks. These 105 × 105 matrices display the t-value in the lower triangle, where each cell reflects a group difference in connectivity between two msICNs. The upper triangle shows the statistical significance using -log(q-value).*sign(t-value) for the discovery set and -log(p-value).*sign(t-value) for the replication set, with non-significant msFNC pairs grayed out. The t-value ranges from -13 to 12. A-C) msFNC group difference matrix between CON vs. PIN-1 in discovery and replication sets and common group differences between them. A) Common group differences of discovery and replication sets. B) Group differences in the discovery set. C) Group differences in the replication set. D-F) msFNC group difference matrix between CON vs. PIN-2 in the discovery and replication sets and common differences between them. D) Common group differences in the discovery and replication sets. E) Group differences in the discovery set. F) Group differences in the replication set. G-I) msFNC group difference matrix between CON vs. PIN-3 in discovery and replication sets and common differences between them. G) Common group differences in the discovery and replication sets. H) Group differences in the discovery set. I) Group differences in the replication set. List of abbreviations – CON: Controls; PIN: Psychosis Imaging Neurosubtype; PIN-1: Psychosis Imaging Neurosubtype-1; PIN-2: Psychosis Imaging Neurosubtype-2; PIN-3: Psychosis Imaging Neurosubtype-3;

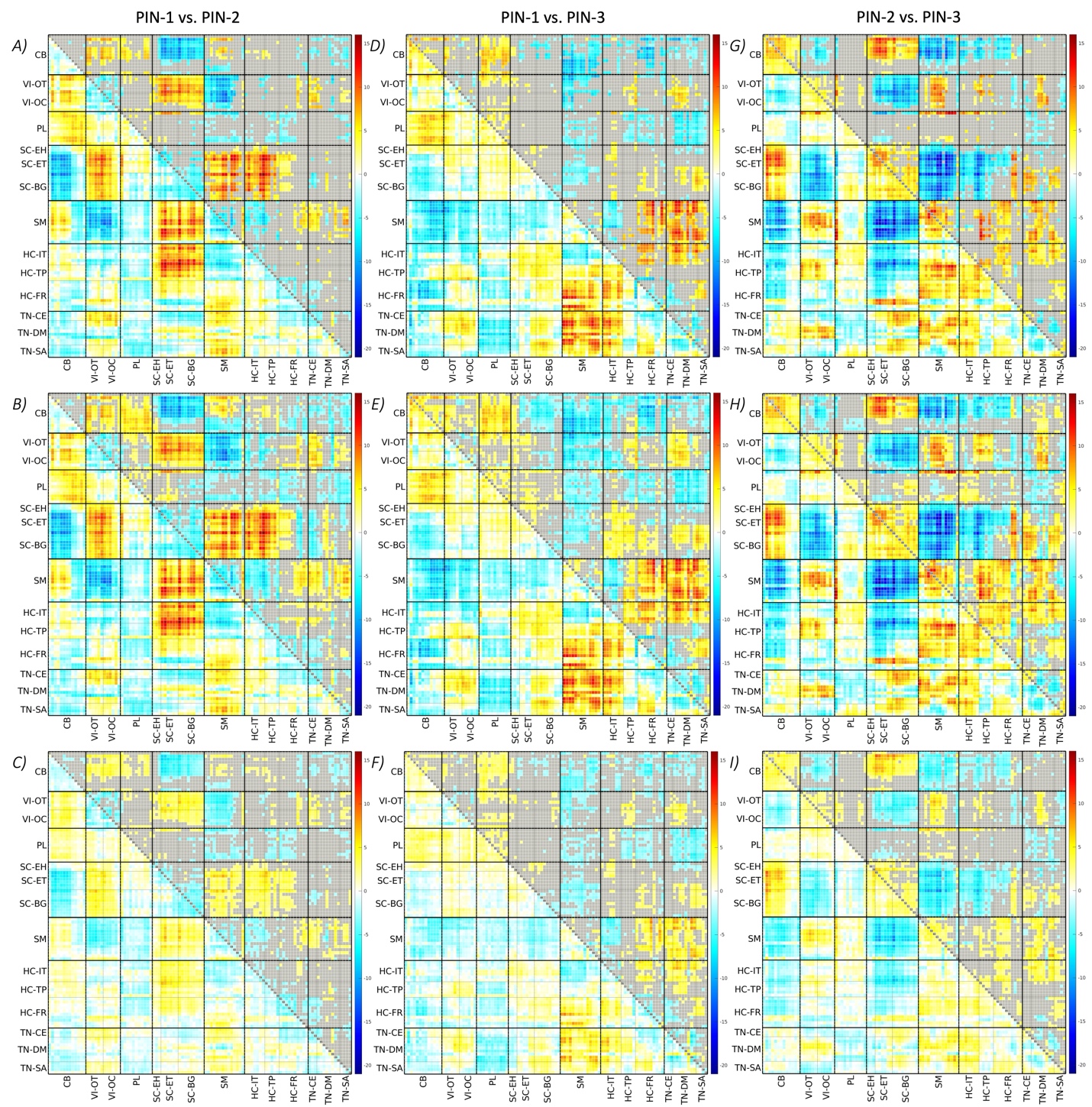

**Supplementary Fig.9**: **Group Differences between Psychosis Imaging Neurosubtypes (PINs) along Multi-scale Functional Network Connectivity (msFNC) in the Discovery and Replication sets.** msFNC represents connectivity between 105 Multi-scale Intrinsic Connectivity Networks (msICNs). Refer Fig.1 and Supplementary Table 2 for additional information on the 105 multi-scale intrinsic connectivity networks. These 105 × 105 matrices display the t-value in the lower triangle, where each cell reflects a group difference in connectivity between two msICNs. The upper triangle shows the statistical significance using -log(q-value).*sign(t-value) for the discovery set and -log(p-value).*sign(t-value) for the replication set, with non-significant msFNC pairs grayed out. The t-value ranges from -21 to 16. A-C) msFNC group difference matrix between PIN-1 vs. PIN-2 in the discovery and replication sets and common differences between them. A) Common group differences in the discovery and replication sets. B) Group differences in the discovery set. C) Group differences in the replication set. D-F) msFNC group difference matrix between PIN-1 vs. PIN-3 in the discovery and replication sets and common differences between them. D) Common group differences in the discovery and replication sets. E) Group differences in the discovery set. F) Group differences in the replication set. G-I) msFNC group difference matrix between PIN-2 vs. PIN-3 in the discovery and replication sets and common differences between them. G) Common group differences in the discovery and replication sets. H) Group differences in the discovery set. I) Group differences in the replication set. List of abbreviations – PIN: Psychosis Imaging Neurosubtype; PIN-1: Psychosis Imaging Neurosubtype-1; PIN-2: Psychosis Imaging Neurosubtype-2; PIN-3: Psychosis Imaging Neurosubtype-3.

**Section 8.4.1. Psychosis Imaging Neurosubtypes - msFNC Variations** **within Functional Domain**

**Visual Domain:** In the VI domain, hypoconnectivity was observed in psychosis PIN-1 in comparison to controls within both OT and OC subdomains (Supplementary Fig.8A). However, PIN-2 and PIN-3 did not show significant connectivity differences compared to controls, except for a small amount of dysconnectivity observed (statistically significant differences but lower msFNC group difference t-values; Supplementary Fig.8D&G). When examining differences among neurosubtypes, PIN-1 displayed decreased connectivity compared to PIN-2 and PIN-3 (Supplementary Fig.9A&D). Additionally, PIN-2 showed increased connectivity compared to PIN-3 within the VI domain (Supplementary Fig.9G).

**Paralimbic Domain:** Dysconnectivity between PINs and controls was observed within the PL domain, though these differences were limited to specific msFNC variations rather than the entire domain (Supplementary Fig.8A,D&G). No significant domain level connectivity differences were found between neurosubtypes and controls. msFNC differences among neurosubtypes also revealed fewer disconnections (Supplementary Fig.9A&G). However, PIN-1 exhibited domain level higher connectivity than PIN-2 within the PL domain (Supplementary Fig.9D).

**Sensorimotor Domain:** In the SM domain, hypoconnectivity was observed in PIN-1 and PIN-3 against controls (Supplementary Fig.8A&G), particularly higher significant differences in the connectivity of ICN 64 (inferior postcentral gyrus) with the entire SM domain. However, PIN-2 and controls did not exhibit any notable connectivity differences (Supplementary Fig.8D). While PIN-1 and PIN-3 exhibited this hypoconnectivity, they exhibited hyperconnectivity compared to controls (Supplementary Fig.8A&G) in the connectivity between ICN 68 (posterior inferior parietal lobe) and the SM domain. But these neurosubtypes PIN-1 and PIN-3 did not show any significant connectivity differences between themselves (Supplementary Fig.9D). PIN-2 however exhibited increased connectivity compared to PIN-3 across the entire SM domain, but lower connectivity along ICN 68 and SM domain connectivity (Supplementary Fig.9G). Conversely, PIN-1 showed lower connectivity compared to PIN-2 across the SM domain, but showed increased connectivity between ICN 68, SM domain connectivity (Supplementary Fig.9A).

**Triple Network Domain:** Dysconnectivity was observed between PINs and controls (Supplementary Fig.8A,D&G) within the TN domain. However, when comparing neurosubtypes, PIN-1 exhibited reduced connectivity in comparison to PIN-3 between ICN 100 (dorsomedial prefrontal cortex in the DM subdomain) and CE subdomain connectivity (Supplementary Fig.9D). Additionally, PIN-2 showed increased connectivity versus PIN-3 within SA subdomain connectivity (ICN 105 - anterior insular, anterior cingulate cortex; Supplementary Fig.9G) and low connectivity against PIN-3 within DM subdomain connectivity (ICN 94, 96 and 97; Supplementary Fig.9G).

**Section 8.4.2. Psychosis Imaging Neurosubtypes - msFNC Variations between Functional Domains**

**Cerebellar - Visual Domains Connectivity:** When analyzing connectivity between CB and VI domains, PIN-1 and PIN-3 exhibited hyperconnectivity compared to controls, particularly in ICN 16, 17, 23, and 24 of the VI domain (Supplementary Fig.8A&G). However, PIN-2 showed hypoconnectivity against controls (Supplementary Fig.8D). Examining psychosis neurosubtypes with each other, PIN-1 displayed increased connectivity compared to PIN-3, particularly with ICN 14 (fusiform gyrus) in the VI domain (OT subdomain), though PIN-1 and PIN-3 had minimal dysconnectivity (Supplementary Fig.9D). PIN-1 also exhibited increased connectivity when compared to PIN-2 (Supplementary Fig.9A), while PIN-2 showed reduced connectivity in comparison to PIN-3 (Supplementary Fig.9G).

**Cerebellar - Paralimbic Domains Connectivity:** PIN-1 displayed hypoconnectivity compared to controls in CB and PL domains (Supplementary Fig.8A), with little dysconnectivity observed between PIN-2, PIN-3 and controls (Supplementary Fig.8D&G). No overall domain connectivity differences were observed between PIN-2 and PIN-3 in PL and CB (Supplementary Fig.9G), except for connectivity differences between ICNs 2 (posterior cerebellum) and 26 (lateral temporal pole). Although, PIN-1 showed overall domain-level increased connectivity in comparison to PIN-3 (Supplementary Fig.9D) than PIN-2 (Supplementary Fig.9A).

**Visual - Paralimbic Domains Connectivity:** Within the connectivity between the VI and PL domains, PIN-1 and controls showed little dysconnectivity (Supplementary Fig.8A), with no significant differences noted between PIN-2, PIN-3 and controls (Supplementary Fig.8D&G). Among the neurosubtypes, VI - PL domains showed no significant differences between PIN-1 and PIN-2, aside from a few msFNC pairs (Supplementary Fig.9A). PIN-1, however, showed higher connectivity than PIN-3 in VI - OT subdomain connectivity with the PL domain (Supplementary Fig.9D). Similarly, PIN-2 exhibited increased connectivity compared to PIN-3 in VI domain connectivity with ICN 26 (lateral temporal pole network; Supplementary Fig.9G).

**Paralimbic - Subcortical Domains Connectivity:** In the connectivity between PL and SC domains no overall differences were found between PINs and controls. However, PIN-1 exhibited hyperconnectivity against controls in ICN 26 (lateral temporal pole network) of the PL domain with SC-ET and BG subdomains (Supplementary Fig.8A). Conversely, PIN-2 showed hypoconnectivity compared with controls in the same networks (Supplementary Fig.8D), while PIN-3 and controls displayed dysconnectivity differences (both hyper- and hypoconnectivity; Supplementary Fig.8G). There were no connectivity differences between PIN-1 and PIN-3 (Supplementary Fig.9D). However, PIN-1 showed increased connectivity versus PIN-2 along ICN 26 with the entire SC domain (Supplementary Fig.9A), while PIN-2 displayed reduced connectivity than PIN-3 (Supplementary Fig.9G).

**Paralimbic - Sensorimotor Domains Connectivity:** In the PL and SM domains, both PIN-1 and PIN-3 displayed hypoconnectivity in comparison to controls, (Supplementary Fig.8A&G), particularly with stronger differences in PIN-3 comparison with controls in ICN 26 connectivity of the PL domain with the entire SM domain. However, PIN-2 showed hyperconnectivity compared to controls (Supplementary Fig.8D). Among the psychosis neurosubtypes, PIN-2 displayed increased connectivity compared to PIN-3 in ICN 26 in the PL domain with the entire SM domain (Supplementary Fig.9G), while PIN-1 showed decreased connectivity when compared to PIN-2 (Supplementary Fig.9A). Both PIN-2 and PIN-3 did not show connectivity differences (Supplementary Fig.9D) in the same ICN 26 connectivity with the SM domain (Supplementary Fig.9D).

**Higher Cognition - Cerebellum Domains Connectivity:** When analyzing HC - CB domain connectivity, PIN-1 showed hypoconnectivity against controls, particularly in ICNs 81, 83, 84, and 86 (overlapping with the frontal and prefrontal cortex) within the HC - FR subdomain (Supplementary Fig.8A). However, PIN-3 exhibited hyperconnectivity versus controls across all HC subdomains (IT, TP, and FR) and the CB domain (Supplementary Fig.8G). PIN-2 and controls had only a few msFNCs showing dysconnectivity (Supplementary Fig.8D). Among neurosubtypes, connectivity differences between HC and CB were also evident. Both PIN-1 and PIN-2 revealed dysconnectivity within these domains (Supplementary Fig.9A), with both exhibiting reduced connectivity when compared to PIN-3. While PIN-1 and PIN-3 showed differences, specifically in HC - FR subdomain connectivity with the CB domain (Supplementary Fig.9D), the connectivity differences between PIN-2 and PIN-3 spanned all ICNs of HC and CB domains (Supplementary Fig.9G).

**Higher Cognition - Visual Domains Connectivity:** When examining HC - VI domain connectivity, PIN-1 and controls displayed minor dysconnectivity (Supplementary Fig.8A), but PIN-1 showed hypoconnectivity between ICN 14 of the VI domain and ICN 81 of the HC - FR subdomain. PIN-3 also exhibited hypoconnectivity compared to controls, particularly in the connectivity of ICNs 76 (left middle temporal gyrus/temporoparietal junction) and 80 (inferior posterior temporal cortex) of the HC - TP subdomain and the VI domain (Supplementary Fig.8D). However, no connectivity differences were present between PIN-2 and controls in these domains (Supplementary Fig.8D). Among neurosubtypes, PIN-1 showed increased connectivity than PIN-2 along ICN 87 in the HC - FR subdomain in connection with the entire VI domain (Supplementary Fig.9A). PIN-1 also displayed increased connectivity compared to PIN-3 along ICN 80 in the HC - TP subdomain with the VI domain (Supplementary Fig.9D). Lastly, PIN-2 exhibited increased connectivity in comparison to PIN-3 in the HC - TP subdomain with the VI domain (Supplementary Fig.9G).

**Higher Cognition - Paralimbic Domains Connectivity:** In the connectivity between the HC and PL domains, PIN-1 displayed hypoconnectivity against controls, particularly along the HC - FR subdomain and PL domain (Supplementary Fig.8A). PIN-1 also showed hypoconnectivity between ICN 14 (fusiform gyrus network) of the VI domain and ICN 81 (right inferior frontal gyrus network) of the HC - FR subdomain. However, PIN-2 exhibited hyperconnectivity versus controls in the connectivity of the HC - IT and TP subdomain and ICN 26 (lateral temporal pole network) of the PL domain (Supplementary Fig.8D). Additionally, PIN-3 showed hypoconnectivity when compared to controls, though this difference was limited to connectivity between the HC - IT subdomain and ICN 26 of the PL domain (Supplementary Fig.8G). When comparing connectivity differences among neurosubtypes, PIN-1 and PIN-2 revealed little dysconnectivity between each other (Supplementary Fig.9A). However, PIN-1 showed higher connectivity compared to PIN-3 along ICN 26 of the PL domain with ICN 72 and 73 of the HC - IT subdomain (Supplementary Fig.9D). PIN-2 also exhibited increased connectivity compared to PIN-3 across all HC subdomains and ICNs 32, 34 and 35 in the PL domain (temporal pole and entorhinal cortex; Supplementary Fig.9G).

**Higher Cognition - Sensorimotor Domains Connectivity:** In the connectivity between HC and SM domains, PIN-2 and controls showed no significant difference (Supplementary Fig.8D). However, PIN-1 revealed hypoconnectivity compared to controls between the HC - IT subdomain and the SM domain and exhibited hyperconnectivity between the HC - FR subdomain and the SM domain (Supplementary Fig.8A). Additionally, PIN-3 showed hypoconnectivity compared to controls between the HC - TP and FR subdomain with the SM domain (Supplementary Fig.8G). Analyzing connectivity differences among neurosubtypes, PIN-1 exhibited lower connectivity compared to PIN-2 and PIN-3 in the connectivity of the HC - IT subdomain and SM domain but displayed higher connectivity compared to both PIN-2 and PIN-3 in the connectivity of the HC - FR subdomain and SM domain (Supplementary Fig.9A&D). PIN-2, on the other hand, revealed increased connectivity than PIN-3 in the connectivity of the HC - TP and FR subdomains with the SM domain (Supplementary Fig.9G).

**Triple Network - Cerebellum Domains Connectivity:** When comparing connectivity differences within the TN and CB domains between controls and PINs, PIN-1 showed hypoconnectivity compared to controls across all TN subdomains (CE, DM and SA) in their connectivity with the CB domain (Supplementary Fig.8A). However, PIN-2 displayed dysconnectivity against controls (Supplementary Fig.8D), and PIN-3 hyperconnectivity compared to controls (Supplementary Fig.8G) across these subdomain connections. The difference between PIN-3 and controls was particularly pronounced in the connectivity between ICN 2 (posterior cerebellum) in the CB domain and the DM subdomain (Supplementary Fig.8G). Regarding connectivity differences among psychosis, PIN-1 showed dysconnectivity compared to PIN-2 (Supplementary Fig.9A), but displayed increased connectivity in comparison to PIN-3, especially in the connectivity between ICN 2 in the CB and the entire TN domain (Supplementary Fig.9D). No significant connectivity differences were observed between PIN-2 and PIN-3 in these domains (Supplementary Fig.9G).

**Triple Network - Visual Domains Connectivity:** In analyzing TN and VI domain connectivity differences between PINs and controls, all comparisons showed dysconnectivity differences (Supplementary Fig.8A,D&G). PIN-1 showed hyperconnectivity compared to controls in the connectivity between ICNs 20 and 24 of the VI - OC subdomain and the TN - CE and DM subdomains (Supplementary Fig.8A). Similarly, PIN-2 showed hyperconnectivity compared to controls in the connectivity of ICNs 15, 21, and 22 of the VI - OT and OC subdomain with ICNs 97 and 99 of the TN - DM subdomain (Supplementary Fig.8D). Among neurosubtypes, PIN-1 displayed increased connectivity compared to both PIN-2 and PIN-3 in the connectivity of the VI domain with TN - CE and DM subdomains (Supplementary Fig.9A&D). However, PIN-1 displayed decreased connectivity than PIN-2 in the connectivity of the VI domain and ICN 99 (dorsal precuneus network) of the TN - DM subdomain (Supplementary Fig.9A). Finally, PIN-2 showed higher connectivity compared to PIN-3 in the VI domain connectivity with the TN - DM subdomain (Supplementary Fig.9G).

**Triple Network - Paralimbic Domains Connectivity:** Along the TN and PL domain connectivity differences among the PINs and controls, PIN-1 showed hypoconnectivity compared to controls within the PL domain’s connectivity with ICN 94 (right inferior parietal lobe/posterior cingulate cortex) of the TN - DM subdomain, as well as between the PL domain and the entire TN - SA subdomain (Supplementary Fig.8A). However, PIN-3 showed dysconnectivity versus controls, specifically hypoconnectivity between the PL domain and ICN 95 (left inferior parietal lobe/posterior cingulate cortex) of the TN - DM subdomain (Supplementary Fig.8G). No connectivity differences were observed between PIN-2 and controls (Supplementary Fig.8D). Along differences between neurosubtypes, PIN-1 and PIN-2 had few dysconnectivity differences (Supplementary Fig.9A). PIN-1 and PIN-3, however, showed differences in overall PL domain connectivity with the entire TN - DM subdomain (Supplementary Fig.9D). Finally, PIN-2 showed reduced connectivity when compared with PIN-3 in the connectivity of ICN 26 with ICN 93, 100, 101, and 103 in the TN - CE, DM and SA subdomains (Supplementary Fig.9G).

**Triple Network - Subcortical Domains Connectivity:** When comparing the TN and SC domain connectivity differences between PINs and controls, PIN-1 demonstrated hyperconnectivity compared to controls in connectivity between the TN - DM subdomain with SC - ET and BG subdomains (Supplementary Fig.8A). Similarly, PIN-2 showed hyperconnectivity in comparison to controls in connectivity between the TN - CE and DM subdomains and the SC - BG subdomain (Supplementary Fig.8D). However, PIN-3 and controls displayed only minor dysconnectivity with each other (Supplementary Fig.8G). Between neurosubtypes, PIN-1 and PIN-2 had few dysconnectivity differences in TN and SC domain connectivity (Supplementary Fig.9A). PIN-1 showed increased connectivity compared to PIN-3 in the connectivity of the TN - DM and SA subdomain with the SC - BG subdomain (Supplementary Fig.9D). However, PIN-2 showed decreased connectivity versus PIN-3 in the connectivity between the TN domain and the SC - ET and BG subdomains, with a more pronounced difference in connectivity between these domains (Supplementary Fig.9G).

**Triple Network - Sensorimotor Domains Connectivity:** Comparing connectivity between PINs and controls along the TN and SM domains, PIN-1 showed hyperconnectivity compared to controls (Supplementary Fig.8A), but PIN-3 showed hypoconnectivity when compared to controls (Supplementary Fig.8G), with differences observed across all ICNs of these domains. No connectivity differences were found between PIN-2 and controls (Supplementary Fig.8D). Among psychosis neurosubtypes, PIN-1 exhibited increased connectivity in comparison to PIN-3 across all ICNs of the TN and SM domains (Supplementary Fig.9D). PIN-2 also displayed higher connectivity than PIN-3, specifically between the TN - CE and DM subdomain with the SM domain (Supplementary Fig.9G). Additionally, PIN-1 showed higher connectivity when compared to PIN-2 in the connectivity between ICNs 91, 95, 100, 103 and 104 of the TN - CE, DM and SA subdomains and the SM domain (Supplementary Fig.9A).

**Triple Network - Higher Cognition Domains Connectivity:** When comparing connectivity between the TN and HC domains, PIN-1, PIN-2 compared to controls, showed little dysconnectivity (Supplementary Fig.8A&D). However, PIN-3 exhibited hypoconnectivity compared to controls in the connectivity between the entire TN domain and the HC - IT subdomain, as well as between the TN - SA subdomain and the HC - FR subdomain (Supplementary Fig.8G). In comparing connectivity differences among neurosubtypes, PIN-1 and PIN-2 displayed no significant differences except for some dysconnectivity in TN and HC connections (Supplementary Fig.9A). PIN-1 showed higher connectivity than PIN-3 between the entire TN domain and the HC - IT subdomain and between the TN - DM, SA subdomains, and the HC - FR subdomain (Supplementary Fig.9D). However, PIN-1 showed reduced connectivity versus PIN-3 in the connectivity between the TN - CE subdomain and the HC - FR subdomain. Additionally, PIN-2 showed increased connectivity compared to PIN-2 across the entire TN domain connectivity with the HC - FR subdomain (Supplementary Fig.9G).

**Section 9. Psychosis First-Degree Relatives Intermediate Neurobiology**

Statistical analyses identified msFNC group differences between psychosis relative participants classified into controls/PINs and the control group from the discovery set. The msFNC differences between controls and relative participants classified into psychosis PIN-1 were highly similar to those observed between controls and psychosis PIN-1 (Supplementary Fig.8A) in the discovery set (rho = 0.86). Similarly, patterns were found for relative participants classified under PIN-2 and PIN-3, with their connectivity differences closely resembling msFNC differences in the discovery set (rho = 0.72 and 0.63 respectively). Additionally, no connectivity differences were found when comparing relative participants identified as controls with the control group from discovery set.

**Section 10. msFNC group differences between Controls and Psychosis participants**

The group analysis of msFNC between control and psychosis participants is shown in Supplementary Fig.4. These results highlight common differences in the upper triangle (Supplementary Fig.4A), with statistically significant findings from the discovery set (Supplementary Fig.4B; q < 0.05) replicated in the replication set (Supplementary Fig.4C; p < 005).

The results indicate that controls showed hyperconnectivity against psychosis group, particularly in the connectivity between CB and SC domains, SC and ICN 88 of HC - FR subdomain, and between SM and HC - TP subdomain (Supplementary Fig.4A). Additionally, hyperconnectivity within the HC domain was observed in the control group. Conversely, controls displayed hypoconnectivity between CB and SM domains, VI and SC - EH subdomain, SC and SM domains, and SC and HC domains (Supplementary Fig.4A).
